# Cell death and growth dynamics shape DNA-replication-based estimates of microbial growth

**DOI:** 10.64898/2026.05.11.723878

**Authors:** Michael Hunter, Hans Ghezzi, Asmita Jain, Jerry He, Carolina Tropini

**Affiliations:** Department of Microbiology and Immunology, University of British Columbia, Vancouver, V6T 1Z3, Canada; School of Biomedical Engineering, University of British Columbia, Vancouver, V6T 1Z3, Canada; Department of Bioinformatics, University of British Columbia, Vancouver, V6T 1Z3, Canada; Humans and the Microbiome Program, CIFAR, Toronto, Canada

## Abstract

A widely used approach for estimating bacterial growth rates is the origin-to-terminus ratio, or peak-to-trough ratio (PTR), which estimates DNA replication activity from origin and terminus copy numbers. Although PTR is theoretically related to bacterial growth rate, it is often compared with, or used to reason about, changes in population size, which reflect the balance between cell division and loss. These quantities may therefore diverge when cell death is appreciable. Here, we used stochastic, cell-based modelling and controlled laboratory experiments to examine how cell death affects the relationship between PTR and changes in population size. We show that transient stress-induced mortality and heterogeneous survival can cause PTR to fail to reflect changes in population size but still reflect bacterial growth rate. We experimentally validated these predictions by exposing *Escherichia coli* to osmotic shock or antibiotics, and measuring change in population size and DNA replication activity. We further show that initial physiological state, lag before replication resumes, and abrupt growth arrest can alter the relationship between PTR and change in population size. In the presence of mortality, lag can make PTR more reflective of changes in population size whereas it decreases its correspondence with bacterial growth rate. Together, these results demonstrate how cell death and changes in replication dynamics can bias the interpretation of PTR-based estimates of population change under controlled laboratory conditions.

## Introduction

The relationship between bacterial chromosome replication and growth has been studied for decades. Most bacteria contain a single circular chromosome that replicates bidirectionally from a defined origin of replication toward a terminus [1]. Consequently, during chromosome replication, regions that have already been replicated exist in two copies, whereas regions yet to be replicated remain in a single copy. Across a population of dividing cells, this produces a genome-wide copy number gradient, where copy number is higher closer to the origin of replication and lower closer to the terminus [2]. Furthermore, it has been shown that the copy number ratio of origins to termini in a population (origin-to-terminus ratio, *R*) is related to the time taken to replicate the chromosome (*C*) and the generation time (*T*) by log_2_(*R*) = *C/T* [2–4]. This therefore establishes a theoretical relationship between origin-to-terminus ratio and bacterial growth rates (defined as the inverse of generation time, 1*/T*).

Peak-to-trough ratio (PTR) was introduced as a computational approach for estimating bacterial growth rates from metagenomic sequencing data [5]. In the last decade, several tools such as iRep [6], GriD [7], DEMIC [8], SMEG [9], and CoPTR [4] have been developed to extend PTR use to situations with incomplete genomes, low coverage, low genomic completion, high genomic redundancy, strain-level specificity, and to improve accuracy. Additionally, it has been shown that origin-to-terminus ratios can be obtained through quantitative PCR (qPCR), with primers targeting the origin and terminus regions, enabling origin-to-terminus ratio measurements without the need for sequencing [10, 11].

The terms origin-to-terminus ratio and peak-to-trough ratio are used somewhat interchangeably throughout the literature. In particular, PTR can refer either to the general approach of using DNA copy number near the origin and terminus, or to the specific computational method originally developed for metagenomic data [5]. Here, we will use the term PTR throughout, even when referring to modeling and qPCR-based approaches, because we believe this is the terminology and usage that appears most commonly in the literature.

PTR-based approaches are commonly benchmarked against bacterial growth rates estimated from measurements of optical density or colony forming units (CFUs) over time [5, 6, 10–13]. Under ideal conditions, such as exponential growth of bacterial populations in nutrient-rich media, these measurements provide reliable estimates of bacterial growth because cell loss is negligible. In general, however, changes in population size represent the net outcome of cell division (i.e., growth) and cell loss (e.g., death), and so PTR is not expected to reliably reflect changes in population size when cell loss is significant [4, 6, 10].

Empirical studies examining the relationship between PTR and changes in population size under more complex conditions have reported mixed results [10, 12, 13]. For example, weak correlations between log_2_(PTR) and rates of population change were reported in marine microbial populations [12]. A subsequent reanalysis using CoPTR found stronger correlations but also identified many negative rates of population change [4]. Although these values were excluded from the analysis, their occurrence indicates that mortality contributed substantially to the observed population dynamics, and likely did so even when the net changes in abundance were positive.

Cell death and other non-idealized conditions may therefore complicate the validation and interpretation of PTR measurements, particularly when they are used to infer changes in population size. To investigate this, we developed an individual-based stochastic model that explicitly tracks DNA replication, cell division, and death in a simulated bacterial population, allowing us to formalize and illustrate how mortality, population heterogeneity, and growth dynamics alter these relationships.

We first analyzed idealized scenarios where replication and death occur at constant rates, establishing baseline conditions under which PTR and changes in population size are closely related. Building on this foundation, we showed that PTR can become uncoupled from changes in population size under common laboratory scenarios in which stress or environmental perturbations alter survival dynamics. Specifically, transient mortality following environmental perturbation, as well as shifts in population composition arising from heterogeneous survival, can cause PTR to reflect bacterial growth rate but fail to accurately capture changes in population size. We further used the model to explore how the initial physiological state of the population, lag before replication resumes, and abrupt, non-lethal growth arrest can influence the relationship between PTR and changes in population size. Together, our computational and experimental results demonstrate how cell death can cause PTR to diverge from changes in population size under controlled laboratory conditions.

## Materials and Methods

### Computational modeling framework

We developed an individual-based stochastic model to explore how variation in DNA replication dynamics and cell death rates affect the relationship between PTR, bacterial growth rate, and changes in population size (Fig. 1). The model simulates a population of bacterial cells that divide and die over time, while explicitly tracking the initiation and progression of successive rounds of genome replication. Each cell is characterized by the number of genome origins and termini it contains, along with the number of ongoing replication rounds and their progression status.

**Figure 1:**
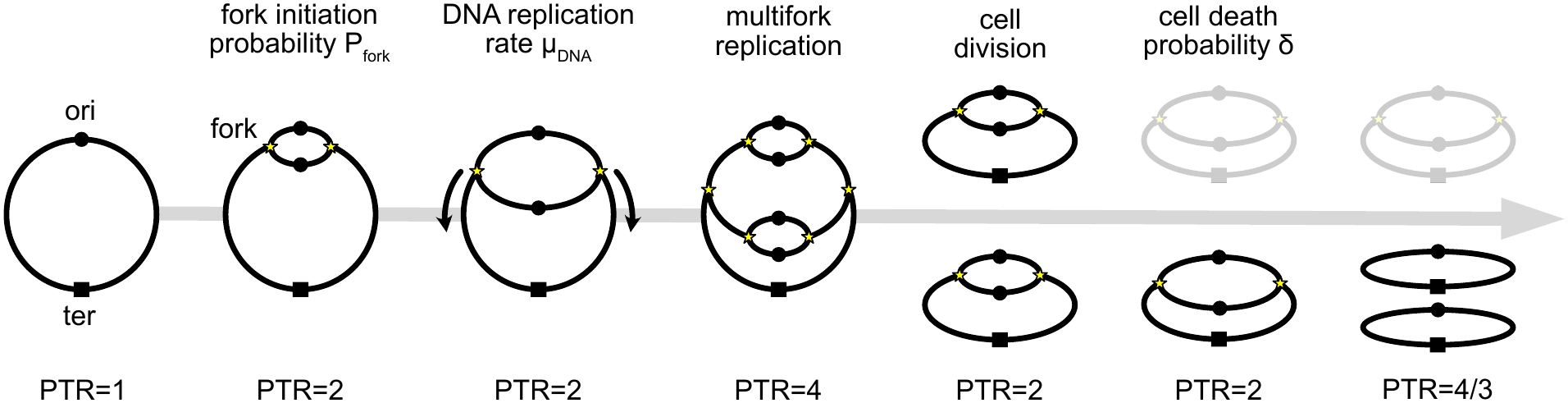
Diagram of computational model framework. Bacterial cells are characterized by the number of origins (ori) and termini (ter) they contain. In each time step, new replication forks are created with probability *P_fork_* and advance towards completion at rate *µ_DNA_*. Multiple forks per cell are permitted and cell division occurs as soon as it contains two complete genome copies. Cell death occurs with probability *δ* in each time step (hereafter referred to as the death rate).

In each simulation time step, new replication forks are initiated with probability *P_fork_*, with replication initiating simultaneously at all existing origins. This assumption does not qualitatively affect our conclusions, as demonstrated by simulations in which replication initiation occurs independently at individual origins (Supplementary Information Section S1). Each replication round progresses towards completion over a fixed *C*-period (i.e., the time taken to replicate the chromosome). Each initiation event doubles the number of origins in the cell, whereas completion of a replication round doubles the number of termini, representing the completion of genome replication. A cell divides as soon as it contains two complete copies of the genome, effectively assuming a *D*-period of zero. Upon cellular division, the number of origins is divided equally between the two daughter cells, which inherit identical replication progression states and subsequently continue growing independently.

Cell death occurs with probability *δ* in each time step; once a cell dies, fork initiation and DNA replication cease immediately, and the dead cell DNA retains the number of origins and termini present at the time of death. Because time proceeds in dimensionless steps rather than continuous units, *δ* also represents the per-step death rate. Thus, *δ* can be interpreted both as the probability of death per cell per step, and as the expected fractional loss of the population per step.

For each simulation, we tracked the numbers of live and dead cells, as well as the PTR calculated from the combined population and from live cells alone. We assume that experimental PTR measurements may include DNA originating from nonviable cells, because such DNA can persist in the environment following cell death for periods ranging from hours or days to weeks or months across laboratory, freshwater, and soil systems, and may therefore be detected alongside DNA from viable cells [14–20]. The combined PTR therefore served as our default measurement, whereas the PTR calculated from live cells alone (PTR*_live_*) reflects the instantaneous bacterial growth state in our simulations. In our primary simulations, we do not account for cells that undergo sudden growth arrest, for example through induction of the stringent response [21–24]; the effects of such growth arrest on PTR are considered separately in Supplementary Information Section S2. Comparing these two PTR estimates thus allowed us to quantify the extent to which DNA from dead cells biases PTR-based estimates of bacterial growth rate. We systematically varied the probabilities of fork initiation, *P_fork_*, the driver of cellular growth in our simulations, and cell death, *δ*, to examine how changes in replication dynamics and mortality shape the relationships between PTR, bacterial growth rate, and changes in population size.

Finally, we assumed a fixed *C*-period of *C* = 100 steps throughout, corresponding to a DNA replication rate of *µ_DNA_* = 1% per step. Consequently, simulation time does not correspond to a fixed physical unit, but is instead defined relative to the *C*-period. For example, if the *C*-period is 100 mins, then one simulation step corresponds to one minute; if the *C*-period is instead 100 hours, then one step corresponds to one hour. This scaling allows the model to represent systems with very different absolute timescales and allows the results to apply broadly to systems with similar qualitative dynamics, regardless of whether they unfold over hours or days.

### Exponential growth simulations

For exponential growth simulations (i.e., simulations with constant rates of fork initiation and death; Fig. 2), simulations were initialized with 10 bacterial cells (each without DNA-replication forks) and run until they either went extinct or surpassed 10^5^, which we set as the carrying capacity. For populations that survived, a per-capita rate of population change (*r*) was estimated by fitting a straight line to the log-transformed population size as a function of time, using data collected after the population exceeded 10^2^. PTR was calculated independently at each time point. For each simulation, the steady-state PTR was then estimated as the mean value across all timepoints after the population surpassed 10^4^, when transient and stochastic effects had largely diminished (Fig. S3). Simulations were run for various *P_fork_* and *δ*.

**Figure 2:**
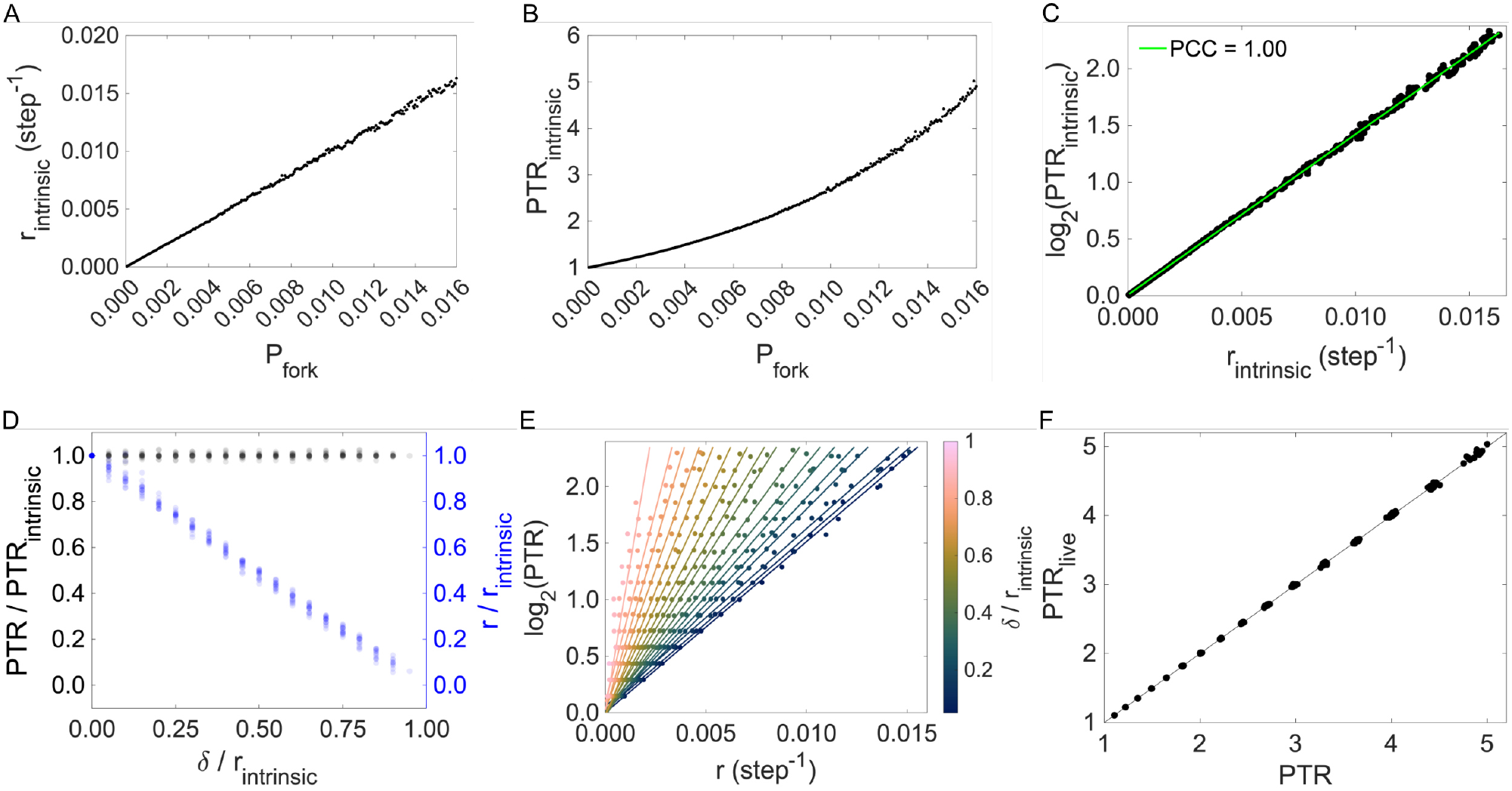
Under constant fork initiation and death probabilities, PTR tracks changes in population size within regimes of fixed relative mortality. (A) Intrinsic – defined as in the absence of cell death – per-capita rate of population change (*r_intrinsic_*), estimated from simulated population size over time (see Methods), as a function of fork probability (*P_fork_*). (B) Intrinsic PTR (PTR*_intrinsic_*) as a function of fork probability. (C) log_2_(PTR*_intrinsic_*) as a function of *r_intrinsic_*. The Pearson correlation coefficient (PCC) equals 1.00. (D) Relative PTR (PTR/PTR*_intrinsic_* – black) and relative per-capita rate of population change (*r*/*r_intrinsic_* – blue) as a function of relative death rate (*δ*/*r_intrinsic_*). (E) log_2_(PTR) as a function of *r*. Colour represents the relative death rate, with linear fits shown for fixed values of the relative death rate (PCC *≈* 1.00 in all cases). (F) PTR measured only from the DNA of live cells (PTR*_live_*) as a function of the PTR measured from all cells. The solid black line indicates a 1:1 relationship.

### Measuring instantaneous per-capita rate of population change from simulation data

In all simulations outside of constant exponential growth, the per-capita rate of population change (*r*) was estimated using a sliding window approach. At each timepoint, a symmetric window spanning 25 points on either side (bounded by the available data) was extracted and a linear fit to the log-transformed population size was performed. The slope of this local fit was recorded as *r* for that time point. Correlation with log_2_(PTR) was then determined using the entire time-series. We did not account for a delay between the two signals when determining the correlations reported in the main text, which has been done in some previous studies [5, 6]. However, this analysis was performed and is discussed in Supplementary Information Section S4.

### Time-dependent death simulations

For time-dependent death simulations (Fig. 3), fork initiation was logistic:

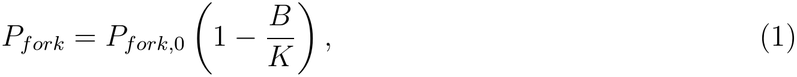

**Figure 3:**
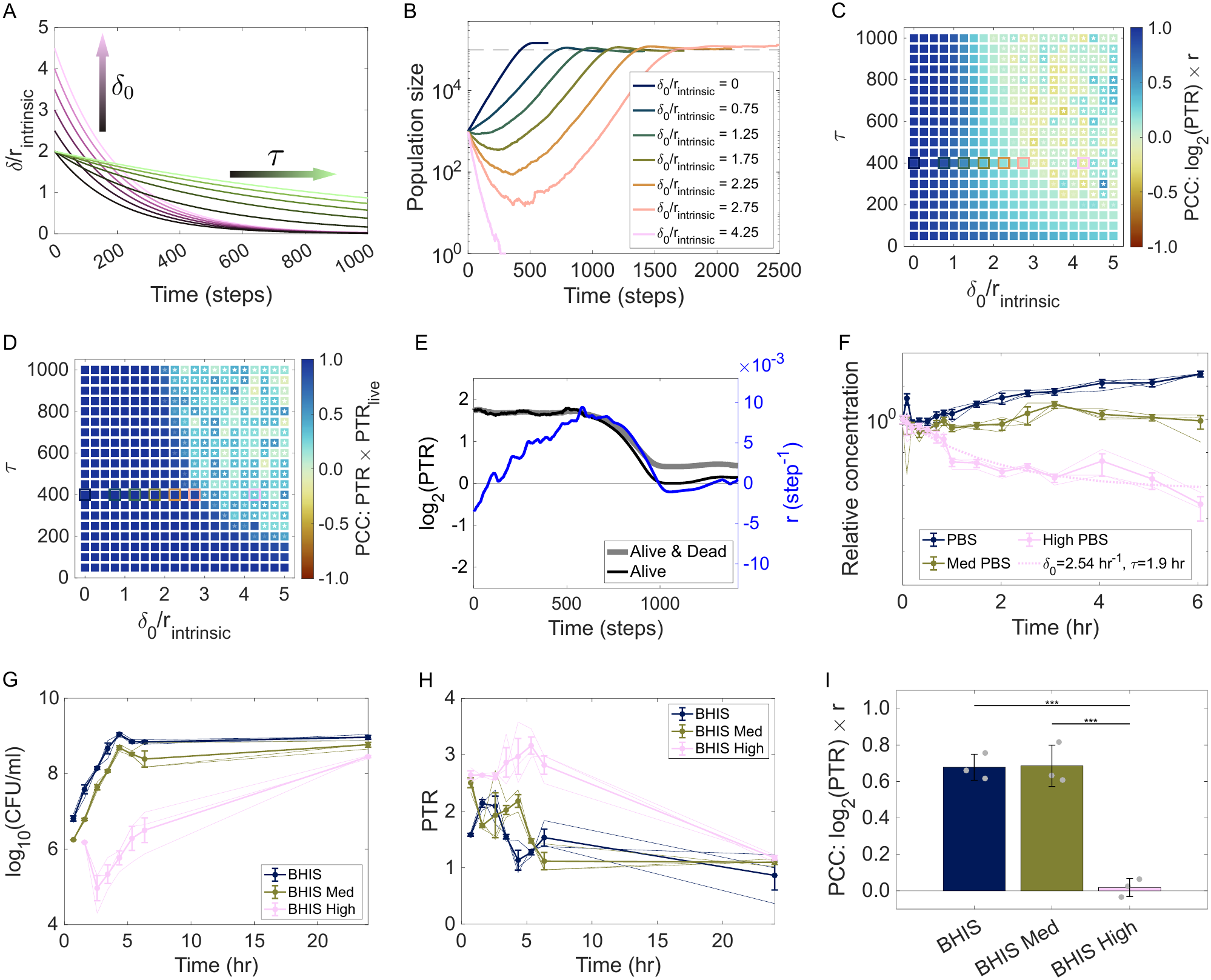
Transient mortality weakens connection between PTR and changes in population size *in silico* (A-E) and *in vitro* in high osmolality (F-I). (A) Relative death rate (*δ*/*r_intrinsic_*) as a function of time - see Eq. 2. The result of increasing initial death rate *δ*_0_ and decay timescale *τ* are shown, relative to *δ*_0_/*r_intrinsic_* = 2 and *τ* = 200 steps. (B) Bacterial population size as a function of time, for various initial relative death rates (*τ* = 400 steps). (C) The Pearson correlation coefficient (PCC) between log_2_(PTR) and *r*, as a function of initial relative death rate and decay timescale. Squares highlight the growth curves shown in (B). (D) The Pearson correlation coefficient (PCC) between PTR measured from all cells and PTR measured from only living cells, as a function of initial relative death rate and decay timescale. Squares highlight the growth curves shown in (B). (E) *r* (blue) and log_2_(PTR) (black/grey) over time for representative simulations with intermediate early mortality (*δ*_0_*/r_intrinsic_* = 1.25, *τ* = 400). (A-E) *P*_*fork*,0_ = 0.012 for all simulations. (F) Survival assays show relative concentration of viable *E. coli* K12 cells as a function of time during incubation in phosphate buffered saline (PBS) supplemented with varying levels of NaCl, following a 1:100 transfer from growth media: baseline PBS (blue, 300 mOsm/kg H_2_O), Med PBS (green, 915 mOsm/kg H_2_O), and High PBS (pink, 1600 mOsm/kg H_2_O). The dotted line shows a fit to the relative concentration data assuming an exponentially decaying death rate of the form given in Eq. 2 – see Section S8 of the Supplementary Information 12 (G) and PTR (H) as a function of time during growth in the rich medium BHIS supplemented with matching NaCl osmolality as (F) over a period of *∼*24 hours. In (F-H), thick lines with markers show the mean of 3 biological replicates (thin lines); error bars show standard error on the mean. (I) The Pearson correlation coefficient (PCC) between log_2_(PTR) and *r* in the different conditions (see Methods). Statistical comparisons performed using one-way ANOVA followed by Tukey’s HSD test. *** *P <* 0.001.

where *P*_*fork*,0_ represents the fork initiation at low density, *B* represents the population of live cells, and *K* represents the carrying capacity. For all the simulations in this section, we used *P*_*fork*,0_ = 0.012 and *K* = 10^5^.

Time-dependent death was modelled using an exponentially decaying death rate, allowing mortality to decline smoothly over time. This functional form was chosen primarily for its relative simplicity and intuitive interpretation, and was also consistent with our experimental data (Fig. 3F). This was intended to capture situations where cells adapt to an initial physiological shock or transient stress condition. Specifically, the death rate *δ* at time *t* is given by

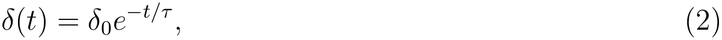

where *δ*_0_ is the initial death rate and *τ* is the death decay timescale (Fig. 3A). A larger *δ*_0_ represents a more severe initial shock, whereas a larger *τ* corresponds to a slower recovery. Together, these parameters control the magnitude and duration of transient mortality following environmental shock.

In these simulations, populations were initialized with 10^3^ exponentially growing cells. This growth state was defined by the PTR in the absence of cell death, hereafter referred to as the *intrinsic* PTR (PTR*_intrinsic_*), obtained from the previous exponential growth simulations. Each cell’s number of replication forks was sampled from a Poisson distribution with mean PTR*_intrinsic_ −* 1, and the initial fork positions were assigned between 0–100% based on the steady-state distribution of fork progressions found in the exponential growth simulations (Fig. S5).

Simulations were run until extinction, attainment of carrying capacity, or – if neither occurred – after a maximum duration of *t_max_* = 3000. When carrying capacity was reached, runs continued until 1.5 times the initial attainment time to ensure entry into a stationary phase. The upper time limit affects situations where the death rate (*δ*) is approximately constant and slightly lower than the intrinsic per-capita rate of population change (*r_intrinsic_*, defined as the per-capita rate of population change in the absence of cell death obtained from the previous exponential growth simulations), which results in quasi-steady states below carrying capacity (Fig. S6).

Results represent averages of five independent simulations (Fig. 3).

### Heterogeneous death simulations

For the heterogeneous population simulations (Fig. 4), the system contained two cell types: one that does not die (referred to as “resistant”), and one that dies at a constant, non-zero rate (referred to as “susceptible”). Fork initiation followed the same logistic form used in the time-dependent death simulations, with shared parameters *P*_*fork*,0_ = 0.012 and *K* = 10^5^. Populations were again initialized with 10^3^ cells in exponential growth.

**Figure 4:**
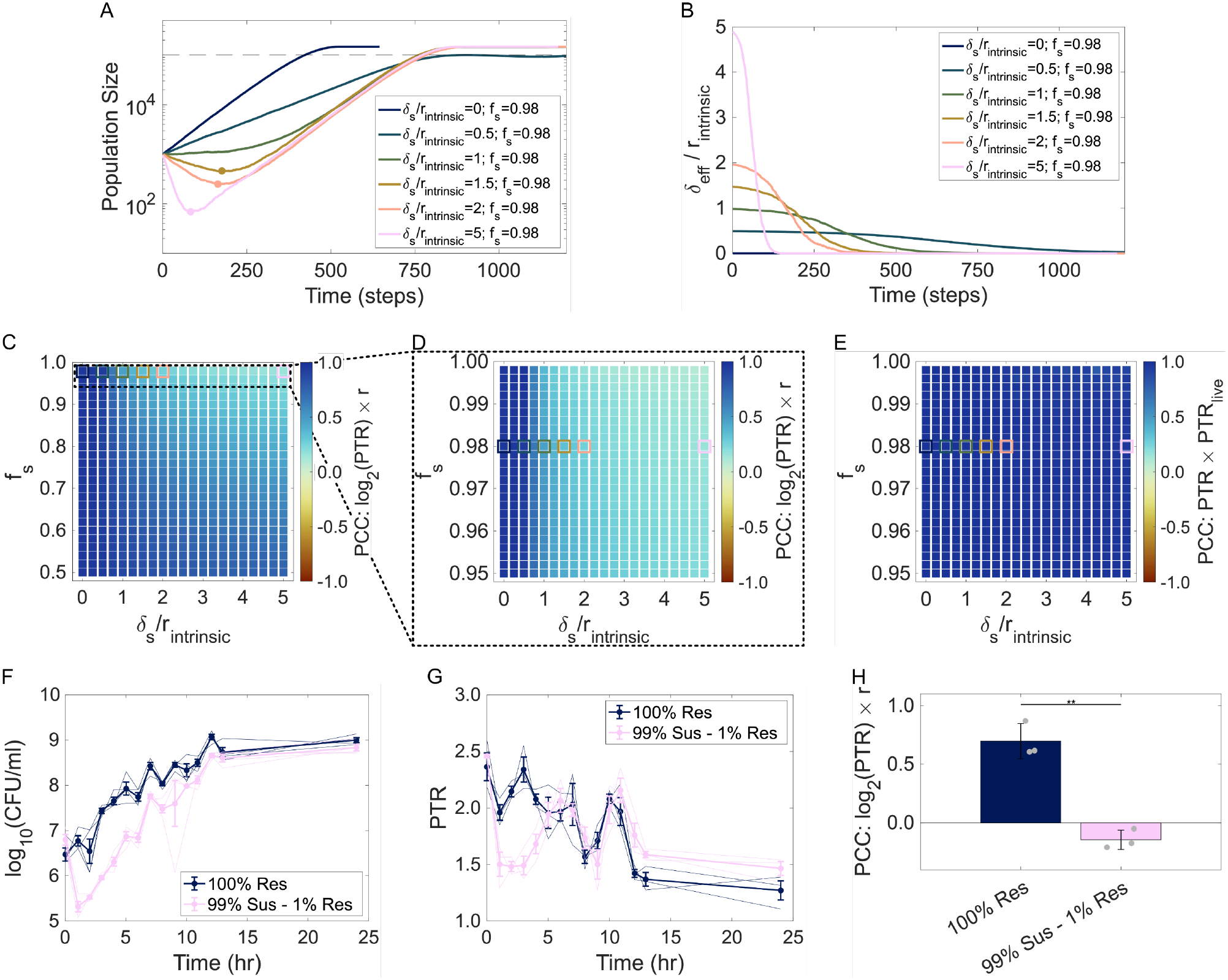
Population heterogeneity generates emergent time-dependent mortality and decouples PTR from changes in population size. (A) Total bacterial population size as a function of time for various death rates of the susceptible fraction *δ_s_*. Initial fraction of susceptible cells *f_s_* is 0.98 and *P*_*fork*,0_ = 0.012 for all curves. Circular markers show the point at which the effective relative death rate – panel (B) – falls below 1. (B) Effective death rate *δ_eff_* of the population as a function of time for each of the growth curves shown in (A). (C, D) The Pearson correlation coefficient (PCC) between log_2_(PTR) and *r*, as a function of the relative death rate and initial fraction of susceptible cells. Squares highlight curves shown in (A) and (B). (E) The Pearson correlation coefficient (PCC) between PTR measured from all cells and PTR measured from only living cells, as a function of the relative death rate and initial fraction of susceptible cells. Squares highlight curves shown in (A) and (B). (F, G) Concentration of viable cells (F) and PTR (G) as a function of time during growth in BHIS supplemented with kanamycin (50 *µ*g/mL) over a period of *∼*24 hours. Blue represents growth of kanamycin-resistant *E. coli* S17-1 only, whereas pink represents kanamycin-susceptible *E. coli* K12 and kanamycin-resistant *E. coli* S17-1 mixed at a 100:1 ratio by cell count. Thick lines with markers show the mean of 3 replicates (thin lines); error bars show standard error on the mean. (H) Pearson correlation coefficient (PCC) between log_2_(PTR) and *r* for the different community compositions (see Methods). Statistical comparison performed using t-test. ** *P <* 0.01.

Only two parameters were varied: the initial fraction of susceptible cells and their death rate. As before, simulations were run until extinction, attainment of carrying capacity, or – if neither occurred – after a maximum duration of *t_max_* = 3000. When carrying capacity was reached, runs continued until 1.5 times the initial attainment time to ensure entry into a stationary phase. Results represent averages over five independent runs (Fig. 4).

### Lag time simulations

For the lag time simulations (Fig. 5), we extended the time-dependent death framework by incorporating a sigmoidal delay in fork initiation and by varying the initial physiological state of the population. Fork initiation followed the same logistic form used in the time-dependent death simulations, but was further multiplied by a “lag term” that modulated the onset of fork initiation – see Eq. 5. The probability of fork initiation at time *t* was therefore

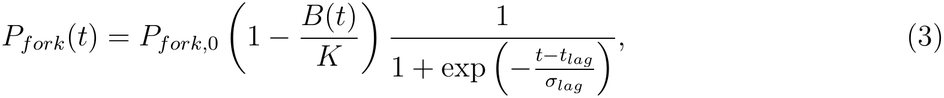

**Figure 5:**
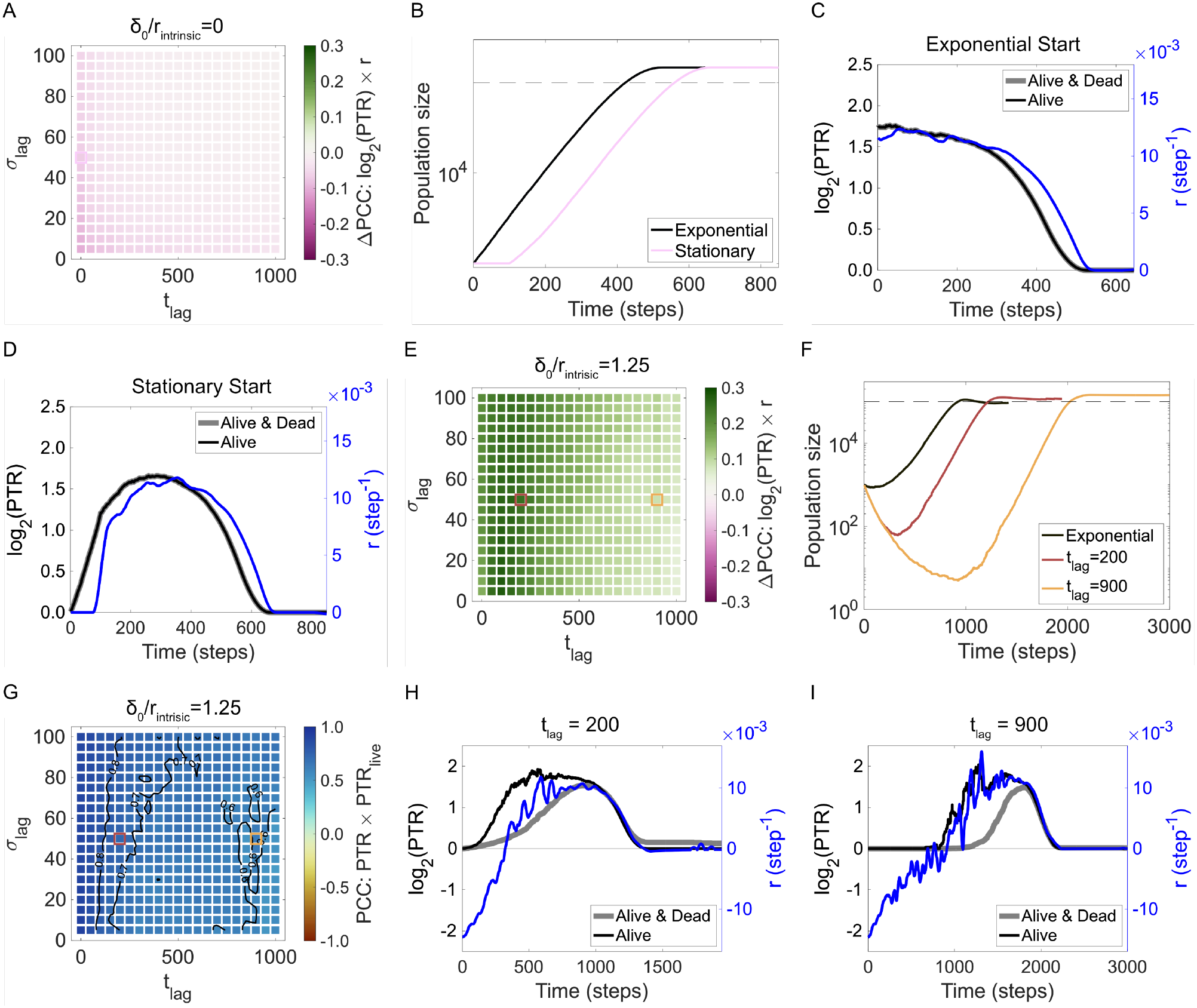
Initial physiological state and lag time modulate the synchrony between PTR and growth. (A) Heatmap showing the difference in correlation between log_2_(PTR) and *r* when simulations are initiated from stationary phase versus exponential phase, in the absence of death (*δ*_0_*/r_intrinsic_* = 0). Black contour line shows ΔPCC = 0.00. The square highlights the parameter combination shown in (B) and (D). (B) Population size over time for simulations initiated from exponential phase or stationary phase with no imposed lag (*t_lag_* = 0, *σ_lag_* = 50). (C, D) *r* (blue) and log_2_(PTR) (black/grey) over time for the exponential-start (C) and stationary-start (D) simulations shown in (B). (E) As in (A), but for simulations with intermediate early mortality (*δ*_0_*/r_intrinsic_* = 1.25, *τ* = 400). The squares highlight the parameter combination shown in (F)–(I). (F) Population size over time for exponential-start simulations, and stationary start simulations with two representative lag times (*t_lag_*= 200 and *t_lag_*= 900, *σ_lag_*= 50). (G) The Pearson correlation coefficient (PCC) between PTR measured from all cells and PTR measured from only living cells. Labelled black contour lines are shown. (H, I) *r* and log_2_(PTR) over time for the *t_lag_* = 200 (H) and *t_lag_* = 900 (I) simulations shown in (F). *P*_*fork*,0_ = 0.012 throughout.

with *P*_*fork*,0_ = 0.012 and *K* = 10^5^ as in the previous sections.

Two factors were varied: the lag parameters (*t_lag_, σ_lag_*) and whether populations were initialized from exponential phase or from stationary phase (i.e., with or without forks, respectively). Exponential-start simulations were identical to the time-dependent-mortality simulations described above (Fig. 3). Lag simulations were performed both without death (*δ*_0_*/r_intrinsic_* = 0) and with intermediate early death (*δ*_0_*/r_intrinsic_* = 1.25, *τ* = 400).

As before, simulations were run until extinction, attainment of carrying capacity, or – if neither occurred – after a maximum duration of *t_max_* = 3000. When carrying capacity was reached, runs continued until 1.5 times the initial attainment time to ensure entry into a stationary phase. Results represent averages over five independent runs (Fig. 5).

### Bacterial strains and culture conditions

#### Escherichia coli

K12 BW25113 (NZ CP009273.1) and *Escherichia coli* S17-1 (CP040667.1) were used. *E. coli* S17-1 was transformed with a plasmid to ensure kanamycin resistance. Briefly, plasmid pJH034 was constructed using NEBuilder HiFi DNA Assembly Master Mix (NEB E2621S). The kanamycin resistance cassette was amplified from the backbone of pK18sB (Addgene #177838) [25] and the R6K origin of replication and origin of transfer (oriT) were amplified from a modified pNBU2 plasmid, pEM105 [26]. Amplification was performed using the manufacturer’s recommended PCR protocol for Phusion High-Fidelity DNA Polymerase (NEB B0518). PCR products were purified in nuclease-free water using the QIAquick PCR Purification Kit (Qiagen 28106). The assembled construct was introduced into TSS-competent *E. coli* S17-1 *λ*pir cells following a previously described transformation protocol [26]. The resulting plasmid was purified by miniprep (Qiagen 27106) and sequence-verified using Oxford Nanopore sequencing (Plasmidsaurus).

Both strains were streaked from glycerol stocks stored at −80°C onto supplemented Brain Heart Infusion (BHIS) agar plates (per liter: 37 g Bacto BHI, 15 g agar for plates, 500 *µ*L vitamin K1 (50 *µ*g in 50 mL 100% EtOH), 1 mL hemin (250 mg in 50 mL 0.1N NaOH), and 1 mL L-cysteine hydrochloride (500 mg/mL in MilliQ water)). Plates were incubated overnight at 37*^◦^*C until single colonies became visible to use in liquid media assays.

### Measuring instantaneous per-capita rate of population change from experimental data

To estimate the per-capita rate of population change (*r*) from experimental CFU/mL measurements, we used the same sliding window approach as for the simulation data. At each timepoint, a symmetric window spanning one (Fig. 3) or two (Fig. 4) points on either side (bounded by the available data) was extracted and a linear fit to the log-transformed population size was performed. The slope of this local fit was recorded as the per-capita rate of population change (*r*) for that time point. Correlation with log_2_(PTR) was then determined using the entire time-series. As before, we did not account for a delay between the two signals when determining the correlations reported in the main text, consistent with some previous studies [5, 6]. However, this analysis was performed and is discussed in Supplementary Information Section S4.

### Replication dynamics under osmotic stress

Biological triplicates of *E. coli* K12 BW25113 single colonies were inoculated into liquid BHIS media and incubated overnight at 37*^◦^*C. Stationary cultures were diluted 1:100, grown until mid-exponential phase at OD_600_ *∼* 0.4, and back-diluted 1:100 into PBS for the death experiment or BHIS media for the growth curves experiment. Experimental media were prepared by supplementing both autoclaved baseline PBS (300 mOsm/kg H_2_O) and BHIS (350 mOsm/kg H_2_O) with NaCl to obtain the “Medium NaCl” (915 mOsm/kg H_2_O) and “High NaCl” (1600 mOsm/kg H_2_O) conditions, followed by sterilization with a 0.2 *µ*m vacuum filter. In the PBS-death experiment, biological triplicates of survival assays in PBS baseline, Medium NaCl, and High NaCl were spot-plated on BHIS plates to obtain CFU/mL every 10 minutes for the first hour since inoculation and then every 30 minutes for the following 5 hours. In the growth curve assay, biological triplicates of cultures in BHIS baseline, Medium NaCl, and High NaCl were spot plated on BHIS plates at 30-45 minute intervals for 6 hours. Bacterial cultures at each timepoint were immediately stored at −20*^◦^*C for DNA extraction and PTR measurements. All BHIS plates were incubated at 37*^◦^*C for 24 hours before counting colonies.

### Replication dynamics in a heterogeneous population

Biological triplicates of *E. coli* K12 BW25113 and transformed *E. coli* S17-1 single colonies were inoculated into liquid BHIS media and incubated overnight at 37*^◦^*C. Stationary cultures were diluted 1:100, grown until mid-exponential phase at OD_600_ *∼* 0.4, and further sub-cultured into BHIS media supplemented with 50 *µ*g/mL kanamycin for 2 experimental conditions: i) only kanamycin-resistant (Res) *E. coli* S17-1, and ii) kanamycin-susceptible (Sus) *E. coli* K12 BW25113 and resistant (Res) *E. coli* S17-1 at a 100:1 ratio. Standard curves to extrapolate CFU/mL from OD_600_ were experimentally established prior to mixing to ensure consistent ratios of both microbes. The concentration of kanamycin used was experimentally validated to be lethal for *E. coli* K12 BW25113 but did not affect transformed *E. coli* S17-1 viability. Conditions (i) and (ii) were spot-plated in biological triplicates on BHIS plates with 50 *µ*g/mL kanamycin and baseline BHIS plates, respectively, for 13 hours at 1-hour intervals. Bacterial cultures were immediately frozen at each timepoint for DNA extraction and PTR measurements. All plates were incubated at 37*^◦^*C for 24 hours before counting colonies.

### Measuring ori:ter ratios

#### Identification of origin and terminus regions and primer design

The genomic position of *oriC* in each *E. coli* strains was identified using ORCA [27], whereas the putative chromosomal terminus position was defined as the point 50% of the genome length from *oriC* [6,28,29]. Primer-BLAST [30] was used with default values to design primers that amplify *∼*150 bp regions located *±* 100 bp before or after the *oriC* and the inferred terminus position in *E. coli* K12 BW25113 (Table S1). Primers were experimentally tested by PCR and gel electrophoresis to also amplify *E. coli* S17-1 and in silico to only target intended regions in both microbes.

#### DNA extraction and quantification

Bacterial cultures from the osmotic stress and heterogeneous population growth curve experiments (see above) were immediately stored at −20*^◦^*C upon collection for simultaneous DNA extraction following the last timepoint. Genomic DNA was extracted using the DNeasy Blood & Tissue Kit (Qiagen) following the manufacturer’s instructions. DNA extraction was performed on 200 *µ*L of the input samples and DNA was eluted in 100 *µ*L of the elution buffer. Genomic DNA concentration was first estimated using a NanoDrop Lite Spectrophotometer (Thermo Fisher Scientific) and then quantified in technical triplicates using the Quant-iT 1× dsDNA HS (High-Sensitivity) Kit (Invitrogen), following the manufacturer’s instructions. Samples with the greatest concentration were used for standard curves using the terminus-region primers in quantitative PCR (qPCR) reactions. Microbial gDNA concentrations measured in nanograms per microliter were converted to genomic copies per microliter using the formula below:

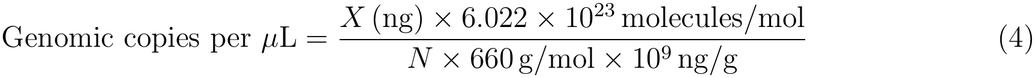

Where *X* is the measured DNA concentration per *µ*L and *N* is the genome length as reported by the NCBI.

In the heterogeneous population experiment, DNA from both *E. coli* strains was pooled at a 1:1 ratio by volume to generate standard curves. The number of genomic copies was calculated by averaging the number of copies of each strain used in the standard curves.

#### Quantification of genomic copies by quantitative polymerase chain reaction

Quantification of *oriC* and terminus region copies was performed with qPCR using a CFX96 Real-Time Module on a C1000 Touch thermocycler (BioRad). Data were collected and analysed using the CFX Manager v.3.1 software (BioRad).

All qPCR reactions performed in the osmotic stress and heterogeneous population experiments contained 12.5 *µ*L of 2× PerfeCTa SYBR Green FastMix (QuantaBio), 10.5 *µ*L of nuclease-free water, 0.5 *µ*L of forward and reverse primers each, and 2 *µ*L of DNA template. Samples were amplified using the following program: 95.0*^◦^*C for 2 minutes, 39 cycles of (95*^◦^*C for 10 seconds, 62*^◦^*C for 30 seconds, 72*^◦^*C for 30 seconds), 72*^◦^*C for 5 minutes, followed by a melting curve from 65*^◦^*C to 95*^◦^*C at 5*^◦^*C increments for 5 seconds each.

Origin-to-terminus ratios (PTR) were then obtained by dividing the genomic copies of *oriC* region by the genomic copies of the inferred terminus region for each technical and biological replicate.

### Materials

All materials used for this project are listed (Table S2).

## Results

### PTR reflects changes in population size under constant relative death rates

We first examined how a constant probability of fork initiation, *P_fork_*, influences *r* (Fig. 2A) and PTR (Fig. 2B) in the absence of cell death – referred to as *intrinsic* per-capita rate of population change (*r_intrinsic_*) and PTR (PTR*_intrinsic_*). As expected, both metrics increased with higher values of *P_fork_*. In this simple case of pure exponential growth, *r* and log_2_(PTR) were perfectly correlated, as expected (Fig. 2C) [4].

Next, we introduced a constant cell death rate and examined its impact on population dynamics. To facilitate comparisons across conditions, we express both the death rate and *r* relative to *r_intrinsic_*. With this scaling, a relative death rate (*δ/r_intrinsic_*) below one corresponds to a population increasing in size, a value of one corresponds to no change in population size, and a value above one corresponds to a population decreasing in size. This normalized representation will also be used throughout the remainder of the manuscript.

As the relative death rate (*δ*/*r_intrinsic_*) increased, *r*/*r_intrinsic_* decreased linearly until it reached zero when the death rate matched *r_intrinsic_* (Fig. 2D). In contrast, PTR remained unchanged, with relative PTR (defined as PTR/PTR*_intrinsic_*) staying constant at 1 across all death rates. This invariance simply reflects our modeling assumptions: DNA replication and cell death are independent processes, so although increased death reduces *r*, it does not alter the rate of fork initiation and thus PTR.

To summarize the relationship between PTR and *r* across all simulated conditions, we plotted log_2_(PTR) against *r* (Fig. 2E) from 400 simulations spanning fork initiation probabilities *P_fork_* = 0.00005 *−* 0.016 and relative death rates *δ/r_intrinsic_* = 0.05 *−* 1. Across this parameter space, *r* ranged from 0 – 0.015 step*^−^*^1^, and PTR ranged from 1 – 5. These PTR values encompass and extend beyond typical physiological measurements, where PTRs are commonly reported from 1 up to *∼*3 [5, 6].

Across this full range, log_2_(PTR) and *r* were not well correlated. Furthermore, this lack of correlation did not arise from a bias in PTR measurements introduced by DNA from dead cells. Because the population remained in steady-state exponential growth and cell death occurred independently of DNA replication, cells that died preserved the same DNA replication states as cells that remained alive. Consequently, PTR measured from all cells, including dead ones, was identical to PTR measured from only living cells (Fig. 2F), indicating that DNA from dead cells did not bias PTR away from the bacterial growth rate. The observed decoupling between PTR and *r* therefore simply reflected the fact that cell death reduced *r* without changing the DNA replication dynamics.

Within regimes of fixed relative mortality, a positive correlation between PTR and *r* was preserved: higher fork probabilities led to faster *r* and higher PTR (Fig. 2E). This can be intuitively understood because, when replication and death occur at constant rates, the population still follows exponential dynamics, just with a reduced *r*, preserving the proportional relationship between PTR and *r*.

### Time-dependent mortality can decouple PTR from changes in population size

Our initial simulations assumed that fork initiation and death probabilities were constant and that populations were initiated with no active replication forks. We next extended our model to better reflect experimental conditions commonly encountered in the lab. In this more realistic scenario, we introduced three key modifications: (i) logistic fork initiation, to capture growth in finite-resource environments where replication slows as populations approach carrying capacity; (ii) initialization from an exponentially growing population, to reflect typical experimental protocols in which cells are sub-cultured from actively dividing cultures; and (iii) a time-dependent death rate, to model transient mortality following environmental perturbations such as stress or antibiotic exposure.

Our choice of a time-dependent death rate was motivated by the fact that bacteria isolated from natural environments are often cultured in laboratory media that differ substantially from their native conditions [31, 32]. This mismatch can cause an initial physiological shock that leads to increased cell death upon transfer, which may decline as the population adapts to the new environment [33–35]. Similarly, time-dependent mortality can arise when lab-adapted strains are exposed to stress conditions, such as altered pH, osmolality, or nutrient availability, where transient death may occur before acclimation [36].

Combining logistic fork initiation, exponential-phase initialization, and time-dependent mortality can produce more complex population dynamics, depending on the choice of parameters (Fig. 3B). For instance, when the initial death rate (*δ*_0_) exceeds *r_intrinsic_*, the population declines in size immediately after transfer. However, because the death rate decays over time, it is possible for the death rate to fall below *r_intrinsic_*before the population goes extinct. At that point, the population begins to increase in size, following a logistic-like trajectory as replication slows with increasing density. Growth continues until the population reaches stationary phase, driven by the saturation of fork initiation due to nutrient limitation.

To evaluate how well PTR reflects changes in population size under these dynamic conditions, we calculated *r* over time using a sliding window approach, and assessed correlation with log_2_(PTR) across the full growth curve (see Methods). We repeated this procedure across a range of parameter combinations, spanning initial relative death rate *δ*_0_*/r_intrinsic_* = 0 *−* 5 and decay timescale *τ* = 50 *−* 1000.

We found that log_2_(PTR) remains strongly correlated with *r* when the initial death rate is low (PCC *≥* 0.90 when *δ*_0_*/r_intrinsic_ ≤* 0.5), regardless of death decay timescale, a regime that approximates logistic growth in which mortality does not substantially perturb population dynamics (Fig. 3C). However, as *δ*_0_ increases, the correlation weakens, indicating that high early mortality can decouple PTR from changes in population size. In contrast, the death decay timescale *τ* had a comparatively modest effect once it exceeded a relatively short threshold (*∼* 100 *−* 300, or 1 *–* 3 genome replications), suggesting that the magnitude of the initial shock is the primary determinant of PTR’s relationship to changes in population size in these scenarios.

The weak correlation between log_2_(PTR) and *r* did not arise from biases in PTR measurements introduced by DNA from dead cells. In all simulations where the bacterial population did not go extinct, PTR measured from all cells was highly correlated with PTR measured only from living cells (Fig. 3D). This is because cells were initialised in an actively replicating state, and most cell death occurred early in the simulation due to the decaying death rate. Therefore, DNA from dead cells predominantly reflected the PTR of cells during exponential growth. Living cells remained in this growing state for most of the growth curve, until the population reached stationary phase, at which point their PTR declined to 1. Consequently, at late time points, DNA from dead cells caused the PTR measured from all cells to be slightly elevated above the PTR of living cells (Fig. 3E), resulting in an overestimation of bacterial growth rate at that time. However, this effect is small because the population had increased by two orders of magnitude from its initial size, meaning that living cells greatly outnumbered dead cells.

To test our modelling predictions experimentally, we used *Escherichia coli* as an extremely well-characterized model organism. PTR-based growth inference was originally validated in *E. coli* under ideal growth conditions [5], making it a convenient system to examine the impact of time-dependent mortality. To induce time-dependent cell death in a controlled manner, we used osmotic shock by supplementing growth media with salt (NaCl). This approach is supported by previous work demonstrating that exposure to osmotic shock with NaCl can induce rapid loss of viability in *E. coli* [37].

To isolate and characterize the death dynamics in absence of growth, cells were grown in rich media (brain-heart infusion supplemented, [BHIS]) to mid-exponential phase and then diluted 100-fold into phosphate-buffered saline (PBS, free of any carbon source) supplemented with NaCl and monitored the population size over time (see Methods). The rationale for using PBS was to decouple growth and death: in nutrient-rich media, cells can simultaneously grow and die, obscuring the underlying death dynamics. In PBS, by contrast, we expect ongoing cell division to be limited due to the lack of nutrients, allowing a clearer measurement of death dynamics. Entry into stationary phase is known to provide cross-protection against osmotic stress [38]. Therefore, to ensure cells retained some level of metabolic activity, we used PBS containing a small carry-over of growth medium (1%, see Methods and Fig. S7). Although this setup minimizes growth, some low-level replication can still occur in base PBS conditions (Fig. 3F). Nevertheless, under high salt conditions, we observed a measurable decrease in viable cell counts consistent with time-dependent mortality (Fig. 3F). However, because mortality was inferred from loss of colony-forming units, these measurements capture loss of culturability, and may include cells that remain viable but fail to form colonies [39].

We then performed growth curves in BHIS containing the same NaCl concentrations as in PBS, periodically measuring population size (Fig. 3G) by spot plating and quantifying PTR (Fig. 3H) using qPCR (see Methods). Under high salt conditions, growth dynamics were characterized by an initial population decline, followed by recovery and growth into stationary phase, closely recapitulating the model predictions (Fig. 3B).

Finally, we calculated *r* using a sliding window approach, and assessed its correlation with log_2_(PTR) over the course of the growth curve (see Methods). In baseline BHIS, *r* and log_2_(PTR) were well correlated; however, under high salt conditions, the relationship broke down, as predicted by the model (Fig. 3I). Because of sampling noise in our experimental setup, we expect correlations *<* 1 even in situations without death (Fig. S8). Overall, these computational and experimental results highlight how time-dependent mortality can transiently decouple PTR from changes in population size, such that PTR may not reliably reflect instantaneous population dynamics during periods of environmental change.

### Population heterogeneity as a source of time-dependent mortality

We next considered how the same decoupling between PTR and changes in population size can emerge from population heterogeneity rather than from explicitly time-dependent mortality. In heterogeneous populations, subpopulations with distinct survival dynamics can change in relative abundance over time, causing the population-level death rate to vary even when the mortality rate of each subpopulation is constant [40]. Here, we used a simple two-population model to illustrate how such emergent changes in population composition influence the relationship between PTR and changes in population size.

Specifically, we modeled a mixed population consisting of two cell types: one that does not die, and one that experiences a constant, non-zero death rate. This setup represents, for instance, a population exposed to a lethal stressor such as an antibiotic, where the majority of cells die, but a small sub-population of resistant mutants survives and continues to grow. More broadly, it captures scenarios in which genetically or phenotypically distinct subpopulations experience different environmental pressures, creating changes in population composition that may not be resolved by PTR measurements.

To explore how sub-population structure affects PTR and growth dynamics, we systematically varied two key parameters: the death rate of the susceptible cell type, and the initial ratio of susceptible to resistant cells. As before, we retained logistic fork initiation and initialized the population from an exponentially growing state.

An illustrative regime occurs when both the death rate of the susceptible population (*δ_s_*) and its initial fraction of the total population (*f_s_*) are high. Under these conditions, the population dynamics resembled those observed when mortality is explicitly time-dependent: the total population initially declined as the susceptible cells rapidly died off, but once they were depleted, the resistant sub-population drove renewed growth (Fig. 4A). Although the death rate of the susceptible fraction in this scenario was constant, the shift in sub-population composition over time created an *effective* time-dependent death rate (*δ_eff_*) at the population level (Fig. 4B), reproducing conditions similar to those examined in the time-dependent-mortality conditions (Fig. 3).

To quantify how PTR reflects growth across the parameter space, we again calculated *r* and assessed its correlation with log_2_(PTR) over time. Across most parameter combinations, log_2_(PTR) and *r* remained well correlated (Fig. 4C). However, when the susceptible population initially dominated, log_2_(PTR) and *r* were correlated only if the death rate of susceptible cells was lower than *r_intrinsic_* (Fig. 4D). As before, this decoupling was not driven by biases from dead cell DNA, because PTR measured from the total population remained strongly correlated with PTR measured exclusively from living cells across the full range of parameters examined (Fig. 4E).

During the population bottleneck and subsequent recovery, PTR and direct measurements of the change in population size provide distinct information about the population dynamics. Although *r* captures the decline of the susceptible population and subsequent expansion of the resistant population, PTR reflects the replication state of the cells contributing to these dynamics. In this case, PTR can remain indicative of bacterial growth rate even during periods in which *r* is negative due to selection against the susceptible subpopulation.

To demonstrate that changes in population composition can occur in a bacterial population and influence PTR-based growth estimates, we experimentally recreated this scenario using two *E. coli* strains: one susceptible and one resistant to the antibiotic kanamycin. Kanamycin was chosen because it is a bactericidal rather than a bacteriostatic antibiotic, and we wanted susceptible cells in our experiment to die rather than simply stop growing, consistent with the situation we modelled – see Ref. [5] for exploration of bacteriostatic antibiotics. We mixed the resistant strain with the susceptible strain at a 1:100 ratio and performed growth curves in BHIS medium supplemented with kanamycin. As in previous experiments, we measured population size (Fig. 4F) over time using spot plating and quantified PTR (Fig. 4G) using qPCR (see Methods). The resulting dynamics closely mirrored the model predictions: the total population initially declined as the susceptible cells were killed by the antibiotic, followed by renewed growth driven by the resistant minority (Fig. 4F). Consistent with the model, this transition in sub-population structure resulted in poor correlation between log_2_(PTR) and *r* over time (Fig. 4H).

### Lag time alters the relationship between PTR, bacterial growth rate, and changes in population size

In the previous sections, we computationally explored scenarios where cell populations were initialized in exponential phase and DNA replication began immediately. Even though these assumptions match common laboratory workflows, they do not fully capture the range of conditions in which such measurements may be used. For instance, in experiments that begin from stationary-phase cultures, cells may experience a lag time before resuming DNA replication.

To explore the effect of lag time, we modulated the fork initiation probability with a sigmoid lag term that controls when fork initiation begins. Specifically, we multiplied the logistic fork initiation probability by [41]

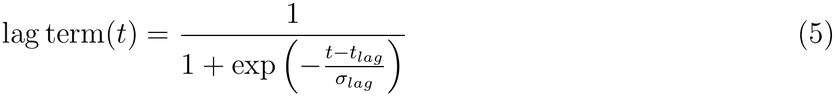

where *t_lag_* controls the time at which fork initiation begins and *σ_lag_* controls how sharply this transition occurs (smaller *σ_lag_* corresponds to a more abrupt transition).

We first considered the case without death (*δ*_0_*/r_intrinsic_* = 0) and compared simulations initiated from stationary phase with those initiated from exponential phase across a range of lag parameters (Fig. 5A). For clarity, the exponential-start simulations correspond exactly to the time-dependent-mortality simulations without a lag term (Fig. 3). For most combinations of lag parameters, the correlations between log_2_(PTR) and *r* in simulations with exponential-start and stationary-start were very similar (ΔPCC *≈* 0.00). However, when fork initiation began immediately (*t_lag_ ≈* 0), starting in stationary phase consistently resulted in slightly lower correlation (Fig. 5A).

The lower correlation in stationary-start simulations with immediate fork initiation occurs because, even without an imposed lag in fork initiation, forks must complete replication before the cells can divide. In exponential-start simulations, new forks were immediately initiated and divisions began without delay (Fig. 5B), producing closely aligned trajectories for *r* and log_2_(PTR) (Fig. 5C). In contrast, populations initialized in stationary phase exhibited a delay of *∼*100 steps before the first cell divisions occurred (Fig. 5B). During this interval, forks continued to be initiated, causing log_2_(PTR) to increase, whereas *r* remained zero (Fig. 5D), thereby slightly reducing the overall correlation between log_2_(PTR) and *r*.

Next, we repeated the analysis in a regime with intermediate early death (*δ*_0_*/r_intrinsic_* = 1.25, *τ* = 400). We chose an intermediate death rate because at higher values of *δ*_0_*/r_intrinsic_*, many combinations of lag parameters led to extinction because the population reached zero before fork initiation resumed, making the correlation undefined over much of the parameter space (Fig. S9). In contrast to the no-death case, starting from stationary phase and adding a fork initiation lag actually improved the correlation between log_2_(PTR) and *r* (Fig. 5E), despite this scenario being characterised by even greater initial death and population decline (Fig. 5F).

The improved correspondence between log_2_(PTR) and *r* came at the expense of agreement between PTR calculated from all cells and PTR calculated from living cells alone (Fig. 5G). Thus, although introducing a lag in fork initiation improves the correspondence between PTR and *r*, it simultaneously reduces its correspondence with bacterial growth rate.

To understand the origin of these effects, we examined representative simulations. In the previous section on time-dependent mortality, we showed that transient mortality weakened the correlation between log_2_(PTR) and *r* because the population initially declined before resuming growth (Fig. 3B). At the same time, PTR measured from all cells remained strongly correlated with PTR measured only from live cells, indicating that dead-cell DNA had little effect on PTR under these conditions (Fig. 3D,E).

By contrast, introducing a lag time delayed the rise in log_2_(PTR), causing *r* and log_2_(PTR) to increase in broadly similar ways (Fig. 5H,I). Although the coupling was not perfect, it avoided the initially flat PTR characteristic of the simulations initialised in exponential phase (Fig. 3E), and therefore yielded substantially higher correlations between log_2_(PTR) and *r*.

At the same time, as the lag time increased, the DNA pool became increasingly dominated by DNA from dead cells prior to replication fork initiation. Because the cells were initialized in stationary phase, they all had a PTR of one. Consequently, although PTR measured from live cells alone increased as soon as replication began, PTR measured from all cells remained lower for longer, owing to the contribution of DNA from dead cells (Fig. 5H,I). This effect intensified with longer lag times, because extended lags resulted in a greater accumulation of dead cell DNA and a correspondingly stronger suppressive effect (Fig. 5H,I). This explains why introducing a lag time reduces the correspondence between PTR measured from all cells and PTR measured from living cells.

## Discussion

It is well established that PTR reflects DNA replication activity rather than changes in population size, and that the two are expected to diverge whenever cell death is non-negligible [4, 6, 10]. However, this understanding is typically stated qualitatively, leaving it unclear how different patterns or levels of death impact the relationship between PTR and changes in population size. This uncertainty complicates the interpretation of PTR measurements in relation to changes in bacterial population size, particularly under conditions where cell death is appreciable.

Here, we used an individual-based stochastic model to characterise and quantify this relationship under conditions commonly encountered in laboratory settings. We show that PTR aligns with changes in population size only when cell mortality is minimal, or scales proportionally with the intrinsic bacterial growth rate. When mortality is instead transient or varies across the population, this correspondence breaks down. This decoupling can arise even when PTR remains representative of the bacterial growth rate, indicating that the divergence stems from population-level dynamics rather than any distortion of PTR itself. In other cases, however, such as when we incorporate a replication lag into our model, DNA persisting from non-viable cells can bias the PTR measurement away from the current replication state of the living population, and thus away from the bacterial growth rate, even though it is more reflective of changes in population size. These results offer a plausible mechanistic account for the discrepancy between the original analysis [12] and subsequent reanalysis [4] of marine microbial communities, where transient mortality and shifts in population composition are both plausible contributors.

Our results also suggest that discordance between PTR and changes in population size could provide an indication that cell death is occurring, even when *r* remains positive. To make use of this, one would simply require paired measurements of PTR and viable population size over time, for example through colony counts over time – measurements that are already collected in studies benchmarking PTR measurements [5, 6, 10–13].

The inverse inference does not hold, however: correlation between PTR and changes in population size does not necessarily indicate that death is negligible. Our simulations that incorporate lag time (Fig. 5) illustrate that death can bias PTR away from the bacterial growth rate even though it is more reflective of changes in population size. Determining whether such discordance reflects cell death therefore requires distinguishing DNA from living and dead cells, for example by measuring PTR using DNA isolated from only living cells. This could be achieved by treating samples with ethidium monoazide (EMA) or propidium monoazide (PMA) prior to DNA extraction [42, 43]. These dyes penetrate the damaged membranes of dead cells and bind DNA, preventing its subsequent amplification and thereby yielding a live-cell-only readout.

Although our work is motivated by conditions commonly encountered during the laboratory validation of PTR-based approaches, the processes captured by our model can also occur in natural microbial communities. Transient mortality following a perturbation (Fig. 3) can occur, such as when the human gut microbiota experiences osmotic stress associated with food intolerances, malabsorption, or laxative use [36]. Likewise, rainfall can trigger pulses of both growth and death in soil microbial communities [44]. Populations in these settings may also resume replication only after a lag (Fig. 5), as dormant or slow-growing taxa are resuscitated once favourable conditions return [45]. The emergent time-dependent death rate that arises from shifts in subpopulation composition in our heterogeneous-population model (Fig. 4) is analogous to a feature of natural microbial communities, in which coexisting taxa differ substantially in their growth rates and stress tolerances [46]. For instance, antibiotic exposure in the human gut microbiota can differentially affect taxa, reshaping community composition during recovery [47]. Although these examples illustrate how the processes considered in our model can arise in natural microbial communities, the extent to which they influence the relationship between PTR and changes in population size under such conditions remains to be established.

Where feasible, PTR measurements in natural settings can be complemented by additional measurements to help distinguish replication dynamics from changes in population size and viability. Longitudinal measurements of absolute cell abundance, for example through culturing or spike-in approaches, can provide an independent measure of population dynamics and identify cases in which changes in abundance are discordant with replication activity inferred from PTR. Measurements of cell viability can further help determine whether cell death contributes to this discordance. In addition, because physiological changes or cell death occurring between sample collection and DNA extraction could alter the measured PTR, sample processing times and conditions should be minimized and standardized. Repeated measurements of the same sample under different processing or storage conditions may also help assess the sensitivity of PTR to these post-sampling effects. Together, these additional measurements may help interpret discrepancies between PTR-based estimates of bacterial replication activity and observed changes in population size.

Our model is simple by design, both so that it is reflective of common laboratory setups that people use to validate PTR-based approaches, and so that the results are easy to interpret. Future work could extend this model to address a broader set of scenarios and assumptions. For instance, we assume a constant rate of DNA replication (and thus a constant C-period), though in reality this can depend on the growth phase [48] and the number of active replication forks [49]. Similarly, we modelled death as independent of growth, but the two are often coupled: for instance, antibiotic-mediated death rates often increase with growth rate in an environment-dependent manner [50]. Finally, our model does not incorporate several features of more complex microbial systems, such as fluctuating conditions, more complex forms of population heterogeneity, and spatial structure, all of which may further influence the relationship between PTR, changes in population size, and bacterial growth rate.

Our results clarify how cell death, population heterogeneity, and lags in the resumption of replication affect the relationship between PTR, bacterial growth rate, and changes in population size under controlled laboratory conditions.

## Acknowledgments

The authors would like to thank: Paula von Sperling, Ian Ghezzi, and Tal Korem for support and useful discussions. This work received support from Advanced Research Computing (ARC) at the University of British Columbia. The authors acknowledge support from Michael Smith Health Research BC Trainee Award (RT-2023-3174, to M.H.), 4-Year Fellowship (to H.G.), Canadian Institute for Advanced Research/Humans and the Microbiome (FL-001253 Appt 3362, to C.T.), Michael Smith Foundation for Health Research Scholar Award (18239, to C.T.), Canada Tier 2 Research Chair, Quantitative Microbiota Biology for Health Applications (CRC-2022-00036, to C.T.), and Canada Foundation for Innovation/Infrastructure Operating Fund (38277), Natural Sciences and Engineering Research Council of Canada (NSERC) Discovery Grants Program (RGPIN-2019-04591, to C.T.). During the preparation of this work, the authors used ChatGPT to edit the manuscript for improved readability. After using this tool, the authors reviewed and edited the content as needed and take full responsibility for the content of the publication. The authors acknowledge that the land we performed this research on is the traditional, ancestral, and unceded territory of the x^w^məθk^w^əýəm (Musqueam) Nation. We encourage others to learn more about the native lands in which they live and work at https://native-land.ca/

## Author contributions

M.H., H.G., and C.T.: conceptualization; M.H. and H.G.: methodology; M.H.: computational modeling; H.G. and A.J.: experimental assays; M.H., H.G., and A.J.: formal analysis, validation, investigation, data curation, and visualization; M.H., H.G., A.J., and C.T.: writing; M.H., H.G., A.J., and J.H.: resources; C.T.: supervision, project administration, and funding acquisition. All authors read and approved the paper prior to submission.

