## Supplemental Information for "Cell death and growth dynamics shape DNA-replication-based estimates of microbial growth"

#### Contents

**S1 Comparing synchronous and asynchronous fork initiation** **1**

**S2 Replication dynamics following non-lethal growth arrest** **2**

**S3 Extracting PTR and $r$ from exponential growth simulations** **2**

**S4 Time delay between PTR and $r$ measurements** **3**

**S5 Steady-state fork distributions** **5**

**S6 Quasi-steady states below the population carrying capacity** **5**

**S7 Cell death in PBS without residual media** **6**

**S8 Derivation of relative population size as a function of time assuming exponentially decaying death rate** **7**

**S9 Sampling noise is predicted to reduce observed correlations even in the absence of death** **8**

**S10 Longer lag times coupled with cell death result in population extinction** **9**

### S1 Comparing synchronous and asynchronous fork initiation

In the model in the main text, we assume that replication initiates simultaneously at all existing origins within a cell, with daughter cells inheriting equal numbers of origins and identical replication progression states. Here, we also consider an alternative model in which replication initiates asynchronously at individual origins, which has been documented in some slow growing bacteria [1]. Upon cellular division, replication forks are randomly distributed between daughter cells.

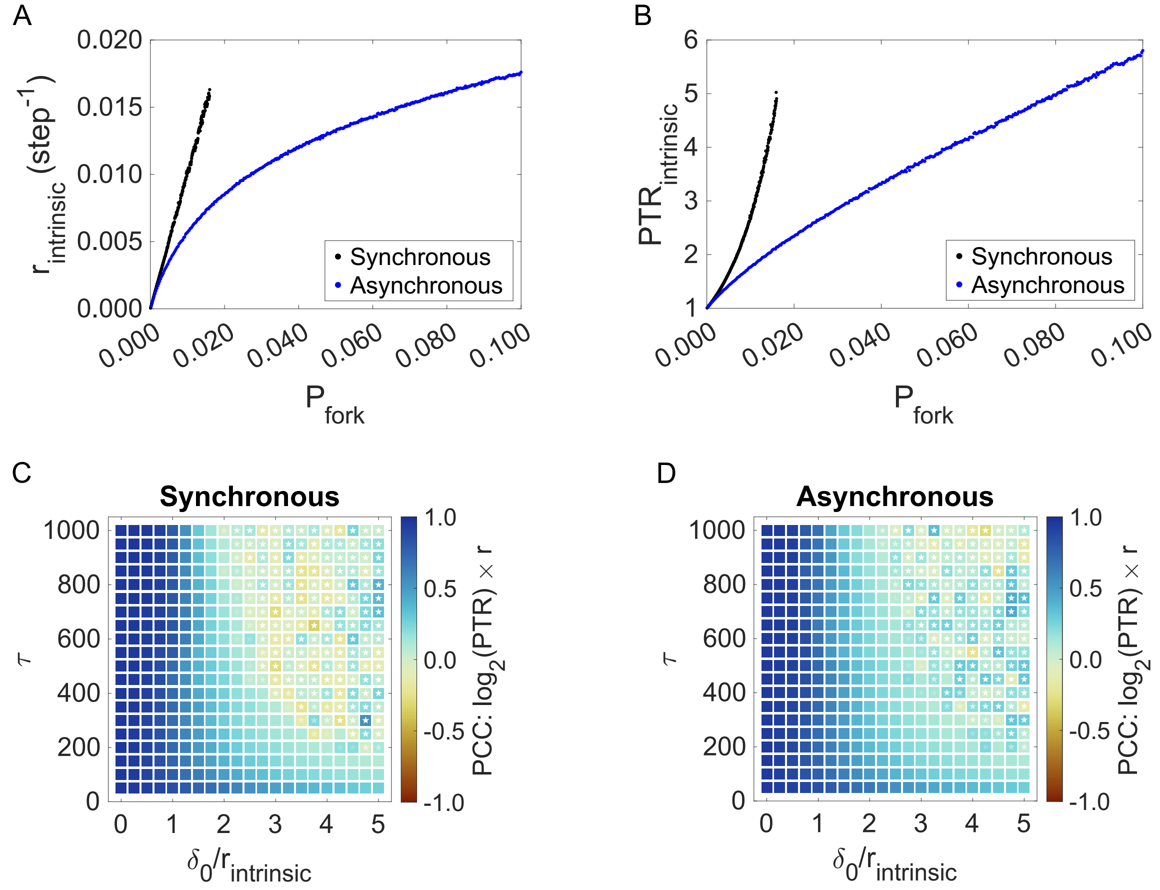


Figure S1: A comparison of synchronous and asynchronous fork initiation. (A) Intrinsic per-capita rate of population change ($r_{intrinsic}$), estimated from simulated population size over time as a function of fork probability ($P_{fork}$) using the two model versions. (B) Intrinsic PTR ($\mathrm{PTR}_{intrinsic}$) as a function of fork probability using the two model versions. (C, D) The Pearson correlation coefficient (PCC) between $\log_2$(PTR) and $r$, as a function of initial relative death rate and decay timescale. In (C) $P_{fork} = 0.012$, and in (D) $P_{fork} = 0.04$.

Although the two models differ quantitatively in the relationship between the probability of fork initiation ($P_{fork}$) and both PTR (Fig. S1A) and the per capita rate of population change ($r$, Fig. S1B), the relationships between PTR, $r$, and the other model parameters remain largely unchanged. In particular, increasing the baseline $P_{fork}$ from 0.012 in the synchronous model to 0.04 in the asynchronous model produces nearly indistinguishable results across the scenarios examined in Figs. 3–5. As an example, Fig. S1C,D compares the correlation between $r$ and $\log_2$(PTR) under explicitly time-dependent mortality for the two models, demonstrating that the results are largely unaffected by the assumption of synchronous versus asynchronous replication initiation.

### S2 Replication dynamics following non-lethal growth arrest

In the model in the main text, we do not account for cells that undergo sudden growth arrest, for example through induction of the stringent response [2, 3, 4, 5]. Here, we present results from simple simulations in which the population was initialised in exponential phase and allowed to grow for a fixed period before growth was abruptly arrested (Fig. S2A).

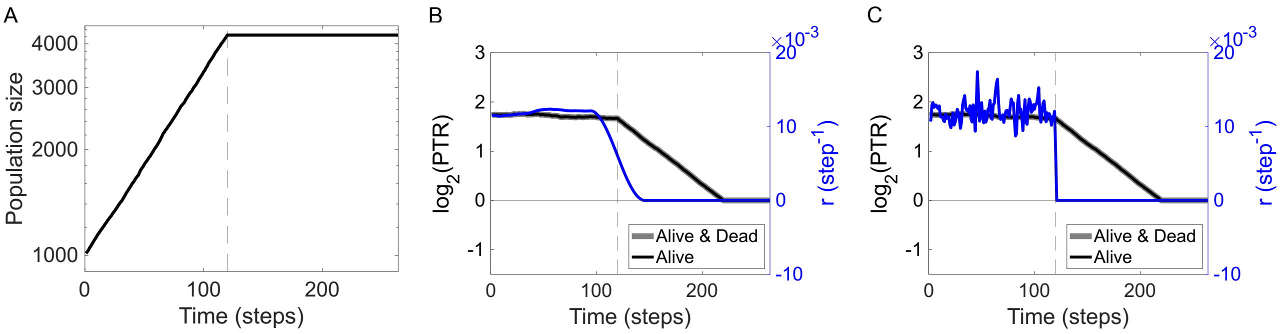


Figure S2: Per-capita rate of population change and PTR respond on different timescales following abrupt growth arrest. (A) Bacterial population size as a function of time. (B,C) Per-capita rate of population change $r$ (blue) and $\log_2$(PTR) (black/grey) over time for the scenario shown in (A). In (B) $n = 25$ neighbouring points on either side were used to measure $r$, whereas in (C) $n = 1$. Throughout, the dashed line indicates the time at which growth was arrested, $P_{fork,0} = 0.012$, and there was no cell death. Plots show the average of 5 simulations.

Following growth arrest, no new replication forks were initiated and no cell division occurred, whereas forks that were already active were allowed to complete replication [2, 4]. As a result, the per-capita rate of population change ($r$) drops quickly to zero upon growth arrest, whereas the PTR gradually declines to 1 over 100 steps (Fig. S2B,C), corresponding to the length of the $C$-period in our model.

Because $r$ is estimated using a sliding-window approach, with $n = 25$ neighbouring points on either side, the measured $r$ does not transition *immediately* to zero following growth arrest (Fig. S2B). When the window is reduced to $n = 1$ neighbouring point on either side, however, the transition is effectively instantaneous, although the resulting estimate for $r$ is correspondingly noisier (Fig. S2C).

### S3 Extracting PTR and $r$ from exponential growth simulations

For the exponential growth simulations presented in the main text (i.e., simulations with constant rates of fork initiation and death) populations were initialized with 10 bacterial cells (each without DNA-replication forks) and run until they either went extinct or surpassed $10^5$ (Fig. S3A). For populations that survived, $r$ was estimated by fitting a straight line to the log-transformed population size as a function of time using data collected after the population exceeded $10^2$ (Fig. S3A). The PTR was calculated as the mean value across all timepoints after the population surpassed $10^4$ (Fig. S3B). These thresholds were selected to reduce measurement noise arising from transient dynamics associated with the initially stationary-phase population, as well as from stochastic fluctuations at small population sizes.

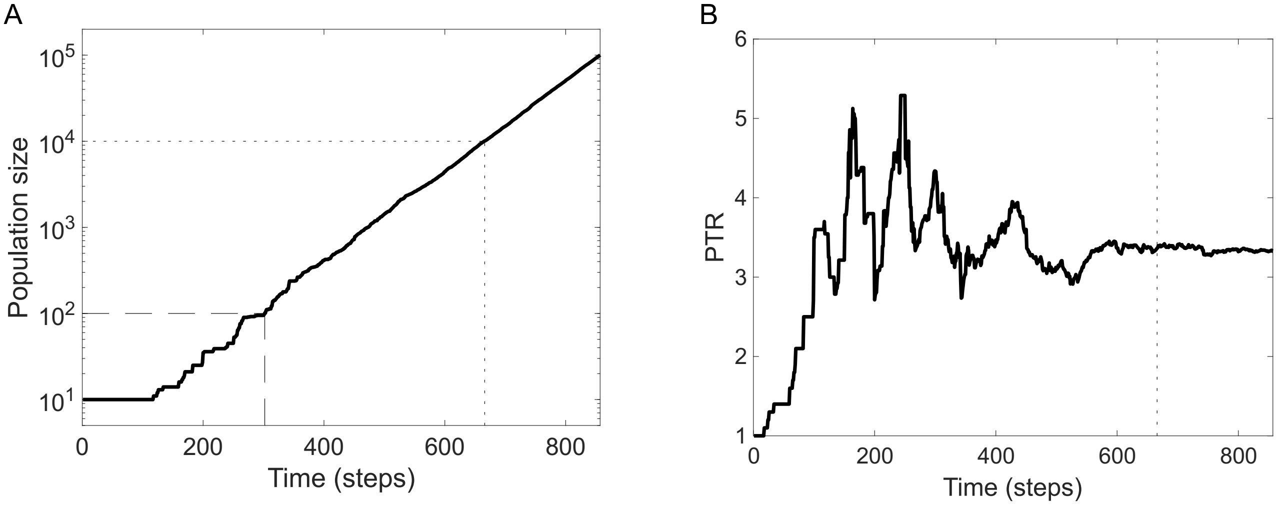


Figure S3: Per-capita rate of population change and steady-state PTR were estimated from different portions of exponential-growth simulations. (A) Bacterial population size as a function of time for a representative exponential growth simulation ($P_{fork} = 0.012$, $\delta = 0$). The dashed line indicates the time at which the population first reaches $10^2$ cells, after which the data were used to estimate $r$. The dotted line indicates the time at which the population first reaches $10^4$ cells, after which the PTR values were averaged to estimate the steady-state PTR. (B) PTR as a function of time for the same simulation. The dotted line again indicates the time at which the population first reaches $10^4$.

### S4 Time delay between PTR and $r$ measurements

The main text presents correlations between $\log_2$(PTR) and $r$ without accounting for a delay between genome replication and observable changes in the population size. The existence of this delay has been described in previous studies comparing PTR with changes in colony forming unit (CFU) counts [6, 7]. In these cases, the delay has been determined empirically by shifting the population size data by varying amounts and identifying the delay that maximises the correlation between $\log_2$(PTR) and $r$. Here, we characterise the effect such a delay would have using the model data from our time-dependent death simulations (main-text Fig. 3).

We first determined the average $r$ and $\log_2$(PTR) across 5 simulations for each combination of the two death parameters (the initial death rate, $\delta_0$, and the death decay timescale, $\tau$). For each parameter combination, we computed the Pearson correlation coefficient (PCC) between $\log_2$(PTR) and $r$ across a range of sample delays, $\Delta t$, and identified the delay that maximised the correlation.

When there was no death, the correlation-maximizing delay was found to be 50 steps, and increased the correlation between $\log_2$(PTR) and $r$ from $\sim$0.98 to $\sim$1 (Fig. S4A,B). Visually, this delay results in close alignment of the two signals (Fig. S4A). Moreover, the 50 step delay is shorter than the 100 step $C$-period in our model, and thus is consistent with the timescale over which the PTR can influence subsequent changes in population size.

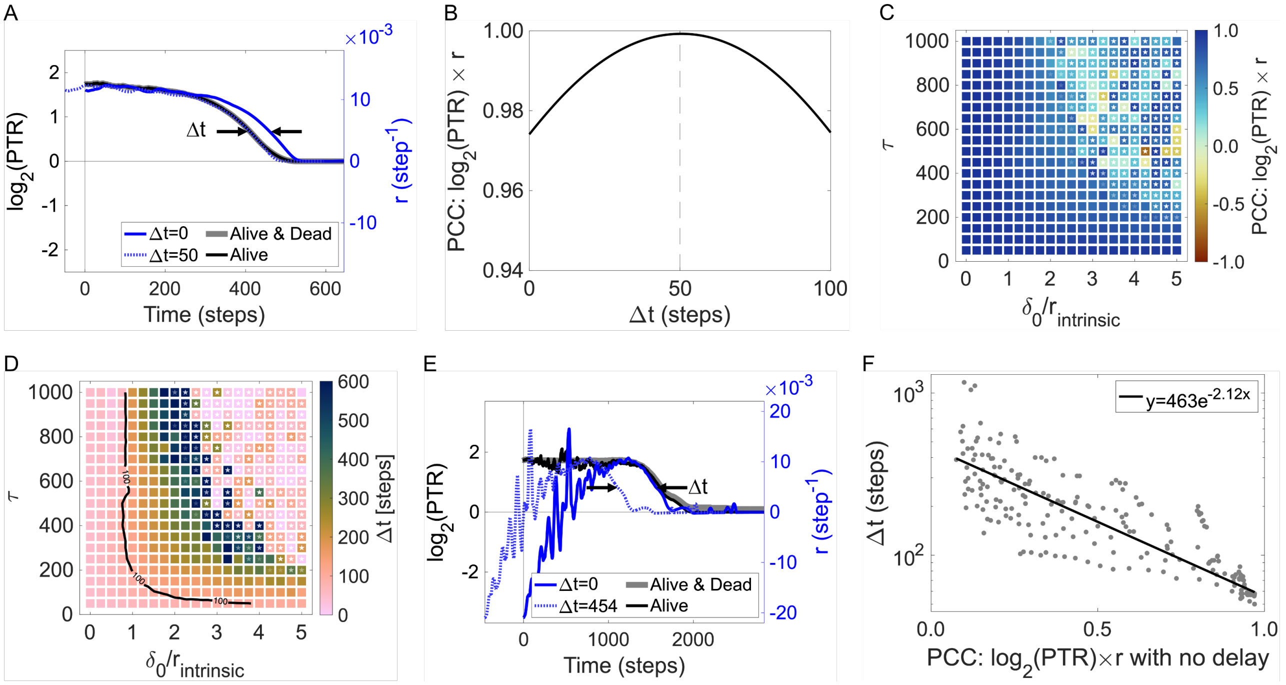


Figure S4: Mortality increases the delay that maximizes the correlation between PTR and $r$. (A) $r$ (blue) and $\log_2$(PTR) (black/grey) over time in the absence of death. $\Delta t$ represents the delay applied to the $r$ data. (B) The Pearson correlation coefficient (PCC) between $\log_2$(PTR) and $r$ as a function of $\Delta t$ for the data shown in (A). The dashed line highlights the value of $\Delta t = 50$ that maximises the PCC. (C, D) The maximised PCC between $\log_2$(PTR) and $r$ (C) and its corresponding delay time $\Delta t$, as a function of initial relative death rate $\delta_0/r_{intrinsic}$ and decay timescale $\tau$. In (D), a black contour highlights values of $\Delta t = 100$. (E) As in (A), but with death parameters $\delta_0/r_{intrinsic} = 2.75$ and $\tau = 400$. (F) The delay time $\Delta t$ that maximises correlation between $\log_2$(PTR) and $r$, as a function of that correlation with no delay (main-text Fig. 3C). $P_{fork,0} = 0.012$ throughout.

Across the full range of death parameters, this procedure yielded substantially higher correlations between $\log_2$(PTR) and $r$ than those reported in the main text (Fig. S4C). However, the correlation-maximizing delay itself also increased with increasing death, often to several hundred time steps (Fig. S4D). Above a relative initial death rate $\delta_0/r_{intrinsic} \sim 1$ (i.e., scenarios where there is an initial decline in population size) the correlation-maximizing delay consistently exceeded 100 steps. This is longer than the 100 step $C$-period in our model, and therefore does not seem consistent with the timescale over which the PTR can influence subsequent changes in population size.

By visually comparing $r$ and $\log_2$(PTR) time series for an example scenario with substantial early death (Fig. S4E), we can see that, unlike the death-free case (Fig. S4A), the increased correlation does not reflect genuine alignment between the two signals. Rather, the large delay simply excludes the majority of the death-affected data from the comparison.

Thus, in the presence of death, determining a correlation-maximizing delay does not simply account for a real delay between genome replication and observable changes in the population size. Instead, it offers an alternative way of quantifying the same death-driven discordance between PTR and $r$ described in the main text. This can be seen by comparing correlation-maximizing delays with the corresponding no-delay coefficients (main-text Fig. 3C; Fig. S4F). For this reason, we report correlations without accounting for a delay in the main text.

### S5 Steady-state fork distributions

In our model, when populations were initialized in an exponentially growing state, the initial progress of the forks were assigned between 0–100% based on the steady-state distribution of fork progressions found in the simple exponential growth simulations (i.e., without death, logistic growth, lag times). Here we show these steady-state distributions for various probabilities of creating new forks $P_{fork}$ – Fig. S5. The distributions were approximately uniform.

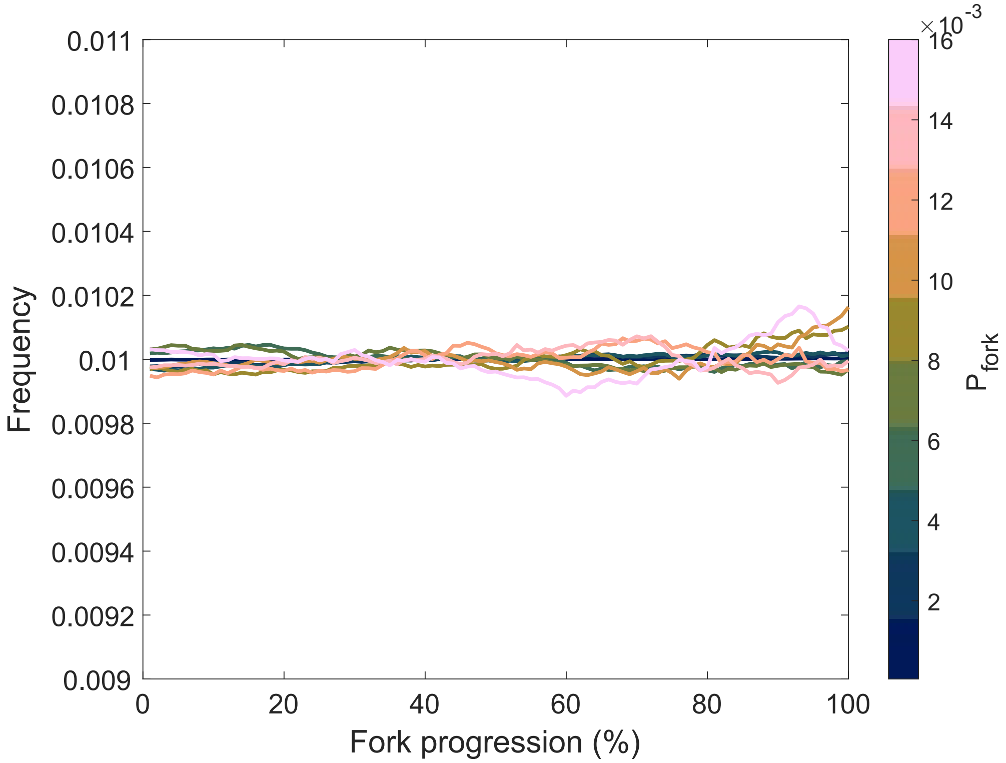


Figure S5: Steady-state distributions of fork progression as a function of fork generation probability $P_{fork}$.

### S6 Quasi-steady states below the population carrying capacity

We imposed an upper time limit of $t_{max} = 3000$ to account for quasi-steady states that arise in the regime $\delta_0/r_{intrinsic} \lesssim 1$ with large $\tau$ (Fig. S6). Due to the fact that fork initiation is logistic in our model, any constant, non-zero relative death rate below one produced a population size below the carrying capacity where the production of new cells is balanced by death, resulting in a steady-state. When the death rate decayed over time, as in our model, this balance was not strictly stable, but could persist as a long-lived quasi-steady state (Fig. S6).

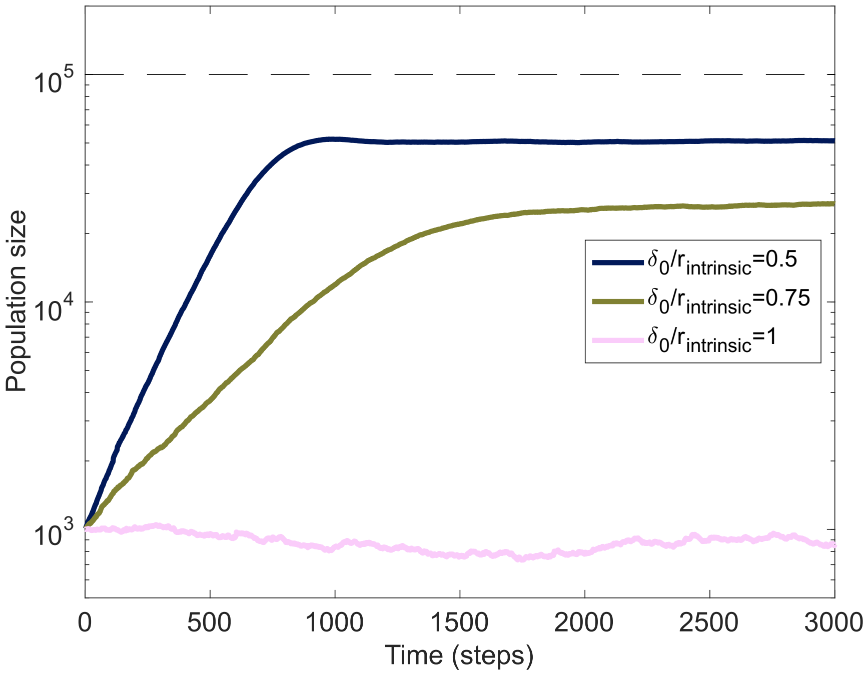


Figure S6: Near-constant mortality can produce long-lived quasi-steady states below carrying capacity. Bacterial population size as a function of time, for various initial relative death rates. $P_{fork} = 0.012$ and $\tau = 100,000$ for all curves. The large decay time was chosen to clearly highlight the quasi-steady states that can occur.

### S7 Cell death in PBS without residual media

In the main text, we isolated and characterized the bacterial death dynamics due to osmotic shock. These experiments were performed by growing bacteria in Brain Heart Infusion Supplemented (BHIS) media until mid-exponential phase ($OD_{600} \sim 0.4$) then diluting 1:100 into phosphate-buffered saline (PBS) supplemented with salt (NaCl) and monitoring the population size over time (Fig. S7A, and Fig. 3F of the main text). Consequently, the PBS contained a residual ($\sim$ 1%) amount of the BHIS media, and some low-level replication still occurred.

To completely remove any growth, we repeated the experiment, but included an additional wash step prior to inoculation into the PBS. The exponential cultures were centrifuged at 8,000 g for 1 min to pellet the cells, the supernatant (media) was removed, and the cells were resuspended in the same volume of PBS. These cultures were diluted 1:100 into the supplemented PBS and the populations monitored as before.

The wash step prevented the populations from growing (Fig. S7B). However, there were also no longer any differences between the different osmotic conditions, with the populations in all three conditions now exhibiting a gradual decline. This is likely due to the fact that nutrient starvation can provide cross protection to osmotic stress [8].

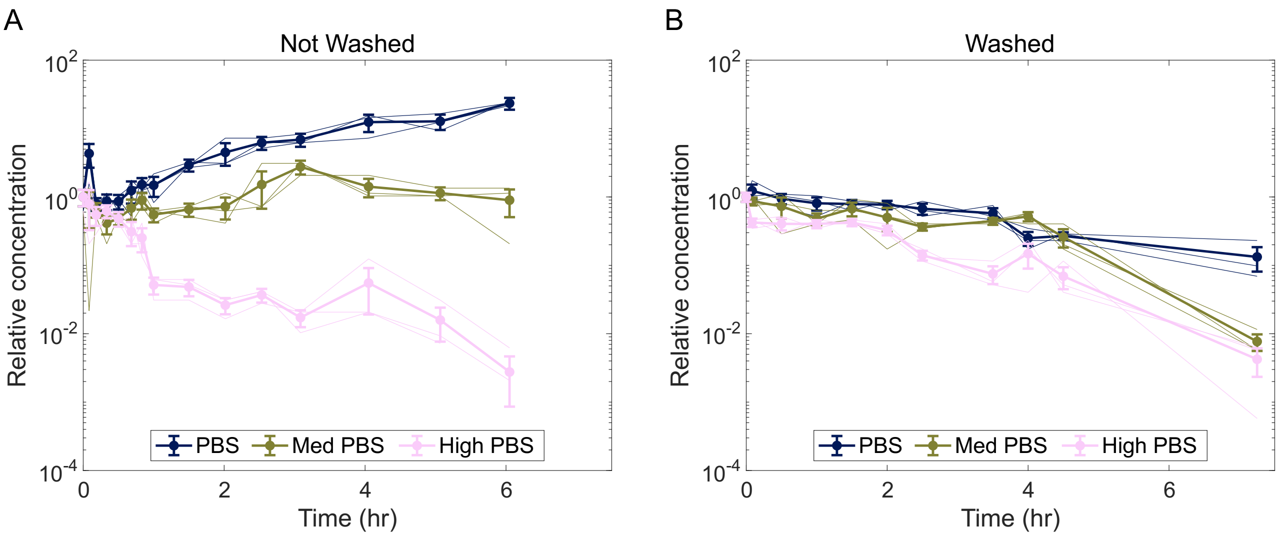


Figure S7: Removing residual growth medium eliminates differences in survival among PBS osmolality conditions. Relative concentration of viable *E. coli* K12 cells as a function of time during incubation in baseline PBS (300 mOsm/kg $H_2O$), Med PBS (915 mOsm/kg $H_2O$), and High PBS (1600 mOsm/kg $H_2O$). Thick lines with markers show the mean of 3 biological replicates (thin lines); error bars show standard error on the mean. In (A) the cells were not washed prior to inoculation into PBS, and so the PBS contains a residual ($\sim$ 1%) amount of the BHIS media. In (B) the cells did undergo a wash step prior to inoculation to remove this residual amount of media.

### S8 Derivation of relative population size as a function of time assuming exponentially decaying death rate

In the main text (and in Section S7) we presented experiments characterising the bacterial death dynamics due to osmotic shock. We wanted to examine whether these observed dynamics were consistent with the exponentially decaying death rate that we implemented in our model, namely

$$\delta(t) = \delta_0 e^{-t/\tau}, \tag{1}$$

where $\delta_0$ is the initial death rate and $\tau$ is the death decay timescale.

To fit this model to our experimental data, we first derive an expression for the population size $N$ as a function of time given the death rate in Eq. 1. We begin with the governing equation for the population size

$$\frac{dN}{dt} = -\delta(t)N, \tag{2}$$

then rearrange, substitute in the expression for $\delta(t)$ given in Eq. 1, and integrate:

$$\frac{1}{N}dN = \delta(t)dt, \tag{3}$$

$$\frac{1}{N}dN = \delta_0 e^{-t/\tau}dt, \tag{4}$$

$$\int \frac{1}{N}dN = \int \delta_0 e^{-t/\tau}dt, \tag{5}$$

$$\ln(N) = \delta_0 \tau e^{-t/\tau} + A, \tag{6}$$

where $A$ represents the constant of integration. This expression can be further rearranged to obtain an expression for $N(t)$ as:

$$N(t) = A' \exp\left[\delta_0 \tau e^{-t/\tau}\right], \tag{7}$$

where $A' = e^A$. To solve for $A'$ we can use the fact that $N(t = 0) = N_0$ (i.e., the fact that the initial population size is $N_0$):

$$N_0 = A' \exp\left[\delta_0 \tau e^{-0/\tau}\right], \tag{8}$$
$$N_0 = A' \exp\left[\delta_0 \tau\right], \tag{9}$$
$$\implies A' = N_0 \exp\left[-\delta_0 \tau\right]. \tag{10}$$

This expression for $A'$ can then be substituted into our previous expression for $N(t)$ to obtain:

$$N(t) = N_0 \exp\left[-\delta_0 \tau\right] \exp\left[\delta_0 \tau e^{-t/\tau}\right], \tag{11}$$
$$= N_0 \exp\left[-\delta_0 \tau + \delta_0 \tau e^{-t/\tau}\right], \tag{12}$$
$$= N_0 \exp\left[-\delta_0 \tau (1 - e^{-t/\tau})\right], \tag{13}$$
$$\implies \frac{N(t)}{N_0} = \exp\left[-\delta_0 \tau (1 - e^{-t/\tau})\right]. \tag{14}$$

The two death parameters ($\delta_0$, $\tau$) can then be determined by fitting our relative concentration data to the above expression for $N(t)/N_0$.

### S9 Sampling noise is predicted to reduce observed correlations even in the absence of death

In the experiments shown in the main text, we observed Pearson correlation coefficients (PCC) between $r$ and $\log_2$(PTR) in the range 0.60–0.80, even in conditions without cell death where we would expect correlations close to 1 (Fig. 3I and Fig. 4H of the main text). We thought that this could likely be explained, at least in part, by sampling noise present in the experiments due to our estimation of population size from colony counts.

To explore this further, we used our model results to simulate experiments many times and look at the distribution of correlation coefficients observed. For a given set of parameters ($P_{fork}$, $\delta_0/r_{intrinsic}$, and $\tau$), the model results were assumed to reflect the underlying population dynamics. We then selected 8 evenly spaced timepoints to match the eight time points sampled in the osmotic-stress experiment (main-text Fig. 3), and simulated a spot plating assay at each of these timepoints. Like in the experiments, the underlying population size $N$ was sequentially reduced (diluted) by a factor of 10. For each dilution $d$, the observed number of colonies $N_{col}$ was drawn from a Poisson distribution with mean $10^{-d}N$. The population was diluted until $1 \leq N_{col} \leq 30$, at which point the estimated population size was calculated as $N_{est} = 10^d N_{col}$.

We then then determined $r$ from the estimated population size at the sampled timepoints using the same sliding window approach employed in the main text. The PCC between $r$ and $\log_2$(PTR) – given by the modelling results – was then determined. We repeated this procedure 1,000 times (i.e., 1,000 simulated experiments) for several different initial relative death rates.

Across all initial relative death rates, the mean PCC obtained from the simulated experiments was lower than the mean obtained from the full modelling results (Fig. S8). The distributions

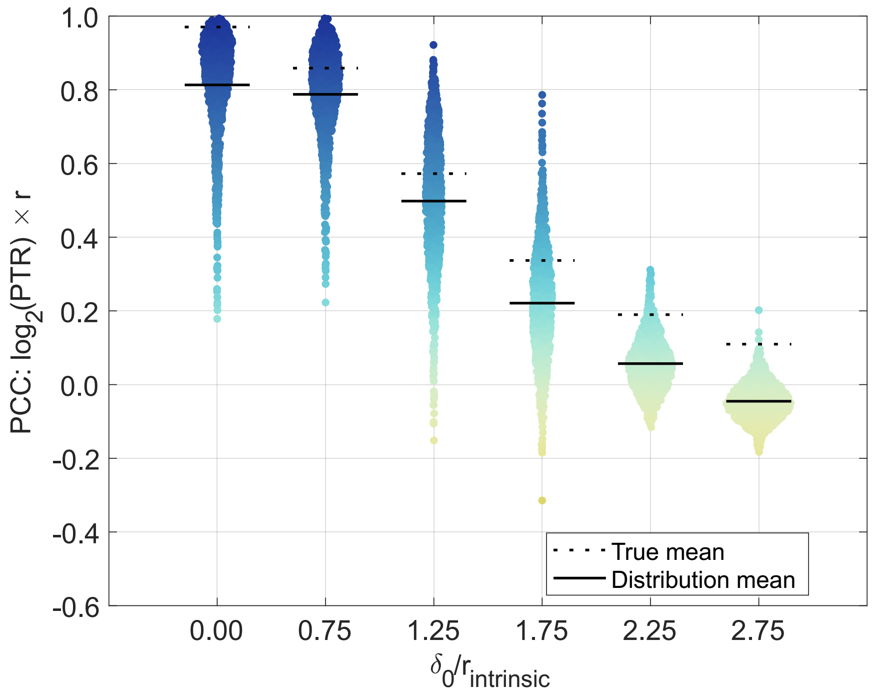


Figure S8: Sampling noise lowers and broadens the observed correlations between $r$ and PTR. Correlations between $r$ and $\log_2$(PTR) observed across multiple simulated experiments (1,000 per initial relative death rate $\delta_0/r_{intrinsic}$). The "True Mean" label refers to the average PCC value determined from the full modelling results.

were also wide, particularly at low and intermediate death rates. This is broadly consistent with our experimental results in the main text, where we observed mean correlations in the range 0.60–0.80 in the absence of death.

It is also worth re-iterating that in our simulated experiments, beyond the small amount of variability between repeat runs of the original model, the only source of variability we accounted for is Poisson sampling noise when estimating the population size. We did not include any error or uncertainty in the measurements of PTR, which almost certainly contributes to variability in our actual experiments. The true experimental distributions of correlations are therefore likely even broader than our simulations suggest.

#### S10 Longer lag times coupled with cell death result in population extinction

In the main text (Fig. 5) we explored how initial physiological state and lag time impacted the relationship between PTR and growth. Here, we performed the same analysis for more initial relative death rates. Consistent with our findings in the main text, when death occurred, starting from stationary phase and including a lag phase generally improved the correlation between $\log_2$(PTR) and $r$, but reduced the correlation between $\log_2$(PTR) and bacterial growth rate (Fig. S9).

Higher initial relative death rates amplified these changes. However, with increasing initial relative death rate, many combinations of lag parameters led to extinction because the population reached zero before fork initiation resumed, making the correlation undefined over much of the

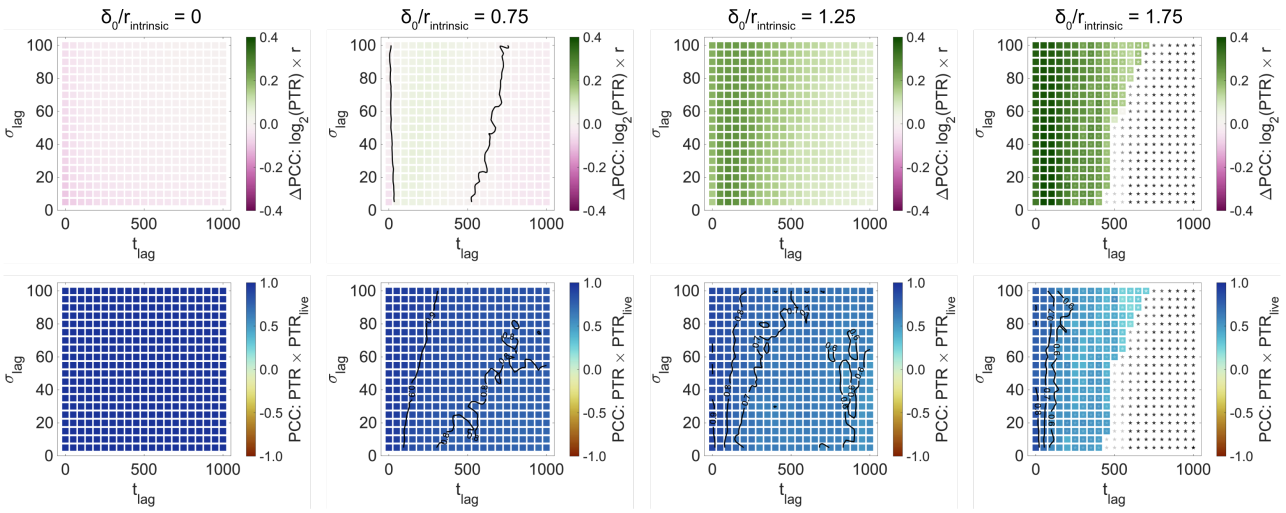


Figure S9: Mortality and lag time jointly determine population survival and the correspondence of PTR with population and bacterial growth rates. (Top Row) Heatmaps showing the difference in Pearson correlation coefficient (PCC) between $\log_2$(PTR) and $r$ when simulations are initiated from stationary phase versus exponential phase. Black contour lines show $\Delta$PCC $= 0$. (Bottom Row) The PCC between PTR measured from all cells and PTR measured from only living cells. Labelled black contour lines are shown. (All) The lag time is defined by the parameters $t_{lag}$, which controls the time at which fork initiation begins, and $\sigma_{lag}$, which controls how sharply this transition occurs (smaller $\sigma_{lag}$ corresponds to a more abrupt transition). Heatmaps are shown for four different initial relative death rates ($\delta_0/r_{intrinsic}$). $\tau = 400$ steps and $P_{fork} = 0.012$ throughout.

parameter space (Fig. S9).

[3] Anurag Kumar Sinha, Anders Løbner-Olesen, and Leise Riber. Bacterial chromosome replication and DNA repair during the stringent response. *Front. Microbiol.*, 11, 2020.

[4] Godefroid Charbon, Belén Mendoza-Chamizo, Christopher Campion, Xiaobo Li, Peter Ruhdal Jensen, Jakob Frimodt-Møller, and Anders Løbner-Olesen. Energy starvation induces a cell cycle arrest in *Escherichia coli* by triggering degradation of the DnaA initiator protein. *Front. Mol. Biosci.*, 8, 2021.

[5] Sophie E. Irving, Naznin R. Choudhury, and Rebecca M. Corrigan. The stringent response and physiological roles of (pp)pGpp in bacteria. *Nat. Rev. Microbiol.*, 19(4):256–271, 2021.

[6] Tal Korem, David Zeevi, Jotham Suez, Adina Weinberger, Tali Avnit-Sagi, Maya Pompan-Lotan, Elad Matot, Ghil Jona, Alon Harmelin, Nadav Cohen, Alexandra Sirota-Madi, Christoph A. Thaiss, Meirav Pevsner-Fischer, Rotem Sorek, Ramnik J. Xavier, Eran Elinav, and Eran Segal. Growth dynamics of gut microbiota in health and disease inferred from single metagenomic samples. *Science*, 349(6252):1101–1106, 2015.

[7] Christopher T. Brown, Matthew R. Olm, Brian C. Thomas, and Jillian F. Banfield. Measurement of bacterial replication rates in microbial communities. *Nat. Biotechnol.*, 34(12):1256–1263, 2016.

[8] Adrienne M. Brauer, Handuo Shi, Petra Anne Levin, and Kerwyn Casey Huang. Physiological and regulatory convergence between osmotic and nutrient stress responses in microbes. *Curr. Opin. Cell Biol.*, 81:102170, 2023.
